# A mechanism that transduces lysosomal damage signals to stress granule formation for cell survival

**DOI:** 10.1101/2024.03.29.587368

**Authors:** Jacob Duran, Suttinee Poolsup, Lee Allers, Monica Rosas Lemus, Qiuying Cheng, Jing Pu, Michelle Salemi, Brett Phinney, Jingyue Jia

## Abstract

Lysosomal damage poses a significant threat to cell survival. Our previous work has reported that lysosomal damage induces stress granule (SG) formation. However, the importance of SG formation in determining cell fate and the precise mechanisms through which lysosomal damage triggers SG formation remains unclear. Here, we show that SG formation is initiated via a novel calcium-dependent pathway and plays a protective role in promoting cell survival in response to lysosomal damage. Mechanistically, we demonstrate that during lysosomal damage, ALIX, a calcium-activated protein, transduces lysosomal damage signals by sensing calcium leakage to induce SG formation by controlling the phosphorylation of eIF2α. ALIX modulates eIF2α phosphorylation by regulating the association between PKR and its activator PACT, with galectin-3 exerting a negative effect on this process. We also found this regulatory event of SG formation occur on damaged lysosomes. Collectively, these investigations reveal novel insights into the precise regulation of SG formation triggered by lysosomal damage, and shed light on the interaction between damaged lysosomes and SGs. Importantly, SG formation is significant for promoting cell survival in the physiological context of lysosomal damage inflicted by SARS-CoV-2 ORF3a, adenovirus infection, Malaria hemozoin, proteopathic tau as well as environmental hazard silica.

## INTRODUCTION

Lysosomes are acidic hydrolase-rich membrane-bound organelles that play a vital role in cellular degradation and signaling(Ballabio et al., 2020; Lamming et al., 2019; Lawrence et al., 2019; C. Yang et al., 2021). Damage to lysosomes can be triggered by numerous physiological and pathological conditions(Nakamura et al., 2021; C. Papadopoulos et al., 2017; H. Yang et al., 2023). These include microbial pathogens(Ghosh et al., 2020; Montespan et al., 2017; Thurston et al., 2012), environmental pollutants(Hornung et al., 2008; Mossman et al., 1998; J. Wang et al., 2017), toxic protein aggregates(Flavin et al., 2017; C. Papadopoulos et al., 2017), endogenous crystals(Hui et al., 2012; Maejima et al., 2013), and many lysosomotropic drugs(Marceau et al., 2012; Pisonero-Vaquero et al., 2017). These agents, along with various others, damage lysosomes, leading to the leakage of acidic contents and the disruption of cellular functions, thereby threatening cell survival(Patra et al., 2023; Saftig et al., 2021; Fengjuan Wang et al., 2018). Lysosomal damage is strongly linked to various human diseases, e.g., cancer, infectious, and neurodegenerative diseases(Amaral et al., 2023; Ballabio et al., 2020; Bonam et al., 2019; Fehrenbacher et al., 2005). Although lysosomal damage is of physiological importance and pathological relevance, understanding of how cells respond to this damage remains largely unknown(C. Papadopoulos et al., 2017).

Cells can detect lysosomal damage through several mechanisms, including the identification of calcium leakage or the exposure of luminal glycan(Aits et al., 2015; Radulovic et al., 2018; Skowyra et al., 2018). Minorly damaged lysosomes can be repaired through multiple cellular systems, including annexins(Ebstrup et al., 2023; Yim et al., 2022), sphingomyelin turnover(Niekamp et al., 2022), microautophagy(Ogura et al., 2023), ER-lysosome lipid transfer(Tan et al., 2022) as well as ESCRT (the endosomal sorting complexes required for transport) machinery(Radulovic et al., 2018; Skowyra et al., 2018). Notably, the protein ALIX (ALG-2-Interacting Protein X), a key ESCRT component, can detect lysosomal damage by sensing calcium release, a function it performs alongside its partner, ALG2 (Apoptosis-Linked Gene-2)(W. Chen et al., 2024; Maki et al., 2016; Sun et al., 2015). Upon detecting such damage, ALIX facilitates the recruitment of other ESCRT components to the site of damage for repair(W. Chen et al., 2024; Radulovic et al., 2018; Skowyra et al., 2018). Severely damaged lysosomes can be removed by selective autophagy(Chauhan et al., 2016; Maejima et al., 2013), noncanonical autophagy(Boyle et al., 2023; Kaur et al., 2023), or lysosomal exocytosis(Wang et al., 2023). Master regulators mTORC1 (mechanistic target of rapamycin complex 1) and AMPK (AMP-activated protein kinase), located on lysosomes(Sancak et al., 2010; C.-S. Zhang et al., 2014), are finely tuned to respond to lysosomal damage, subsequently activating downstream processes e.g., autophagy and lysosomal biogenesis(Jia et al., 2018; Jia et al., 2020a, b; Jia et al., 2020c).These mechanisms collectively safeguard lysosomal quality, maintaining cellular homeostasis(Jia et al., 2020d).

Recently, we reported that lysosomal damage induces the formation of stress granules (SGs)(Jia et al., 2022). SGs are membrane-less organelles identified as ribonucleoprotein condensates that are believed to serve as protective responses in cells under adverse conditions(Ivanov et al., 2019; McCormick et al., 2017; Riggs et al., 2020). Consequently, dysfunctional SGs have been implicated in various human diseases e.g., neurodegenerative and infectious diseases(Advani et al., 2020; Protter et al., 2016; Fei Wang et al., 2020). SG formation is triggered by specific kinases, such as PKR (Protein Kinase R), that sense various stress stimuli, leading to the phosphorylation of eIF2α (eukaryotic translation initiation factor 2)(N.L. Kedersha et al., 1999; Srivastava et al., 1998). Phosphorylated eIF2α (p-eIF2α) halts global translation, resulting in the accumulation of untranslated mRNA(Jackson et al., 2010). Simultaneously, it promotes the selective expression of stress response proteins, a process known as the integrated stress response(Costa-Mattioli et al., 2020; Pakos-Zebrucka et al., 2016). SG formation can also occur through mTORC1-mediated translational shutdown, independent of p-eIF2α(Emara et al., 2012; Fujimura et al., 2012; McCormick et al., 2017). RNA-binding proteins G3BP1/2 (GAP SH3 Domain-Binding Protein 1/2) detect untranslated mRNA and collectively initiate SG formation through an RNA-protein network, driven by liquid-liquid phase separation(Hyman et al., 2014; Ivanov et al., 2019).

Despite the extensive knowledge of SG composition and dynamics, our understanding of the functional consequences of SG formation remains limited(Riggs et al., 2020). Significantly, SG formation has often been investigated under non-physiological conditions such as arsenic stress or heat shock(Jain et al., 2016; Sidrauski et al., 2015; Turakhiya et al., 2018; Verma et al., 2021; P. Yang et al., 2020). Our study(Jia et al., 2022) which originally revealed lysosomal damage as a critical internal physiological trigger for SGs, underscores the need to better understand the nature of SG formation in disease contexts. Additionally, this new connection between damaged lysosomes and SGs provides a novel perspective on the interaction between membrane-bound and membrane-less organelles(Zhao et al., 2020). For example, recent research suggests that SGs have the ability to plug and stabilize damaged lysosomes(Bussi et al., 2023). However, the precise regulation of SG formation in response to lysosomal damage and its consequential impact on cell fate remains largely unexplored.

In this study, we employed unbiased approaches to investigate how lysosomal damage signals are transduced to induce stress granule formation and to elucidate the cytoprotective role of SG formation in promoting cell survival against lysosomal damage. Our findings revealed a novel function of ALIX, which senses calcium release from damaged lysosomes, in controlling the phosphorylation of eIF2α through PKR and its activator on damaged lysosomes, thereby initiating SG formation. This process is critical for cell survival in response to lysosomal damage caused by chemical, pathological and physical agents including SARS-CoV-2 ORF3a, adenovirus, Malaria hemozoin, proteopathic tau and silica. In conclusion, our study uncovers a calcium-dependent signaling mechanism that transmits lysosomal damage signals to induce SG formation and reveals the cytoprotective role of SG formation in response to lysosomal damage caused by diverse agents.

## RESULTS

### Stress granule formation promotes cell survival in response to lysosomal damage

How does SG formation affect cell fate during lysosomal damage? We utilized SG deficient U2OS cells (the human osteosarcoma epithelial cell line) lacking both G3BP1 and G3BP2 genetically (ΔΔG3BP1/2)(Nancy Kedersha et al., 2016), which are essential factors for SG formation(Guillén-Boixet et al., 2020; Nancy Kedersha et al., 2016; P. Yang et al., 2020) (Fig. S1A). We quantified the number of SGs using the canonical SG marker polyA RNA(Ivanov et al., 2019) via high-content microcopy (HCM) (Fig. S1B) and verified the depletion of SG formation in ΔΔG3BP1/2 cells when exposed to the lysosome-specific damaging agent L-leucyl-L-leucine methyl ester (LLOMe)(Jia et al., 2022; Tan et al., 2022; Thiele et al., 1990) (Fig. S1B). A propidium iodide (PI) uptake assay measuring plasma membrane integrity(Crowley et al., 2016; Liu et al., 2023) was adapted to quantify cell survival during lysosomal damage using HCM. We found significant cell death upon LLOMe treatment in ΔΔG3BP1/2 cells compared to wildtype (WT) U2OS cells (Fig. 1A). This is additionally confirmed by using lactate dehydrogenase (LDH) release assay measuring a non-specific leak from cells(Chan et al., 2013; Kumar et al., 2018) (Fig. 1B). Further, we pharmacologically blocked SG assembly through the use of cycloheximide which freezes ribosomes on translating mRNAs and reduces the accumulation of free untranslated mRNA(Freibaum et al., 2021; N. Kedersha et al., 2000). Consistent with previous reports(Bussi et al., 2023; Jia et al., 2022), cycloheximide treatment inhibited SG formation in U2OS cells, as evidenced by the absence of G3BP1 puncta following LLOMe treatment (Fig. S1C). This suppression of SG formation led to reduced cell survival, as indicated by increased LDH release in the face of lysosomal damage (Fig. S1D). Previously we reported that LLOMe treatment induced phosphorylation of eIF2α(Jia et al., 2022), a critical signal for SG formation(Ivanov et al., 2019; N. Kedersha et al., 2000). The small molecule ISRIB (integrated stress response inhibitor) can also act as a SG inhibitor, effectively counteracting the downstream effects of eIF2α phosphorylation, such as ATF4 (Activating transcription factor 4) expression(Rabouw et al., 2019; Sidrauski et al., 2015). We prevented SG formation using ISRIB upon lysosomal damage (Fig. S1E) and observed a corresponding reduction in ATF4 expression levels in THP-1 cells (the human monocytic cell line) (Fig. S1F). The prevention of SG formation by ISRIB also caused a decrease in cell survival in THP-1 cells (Fig. S1G).

**Figure 1.**
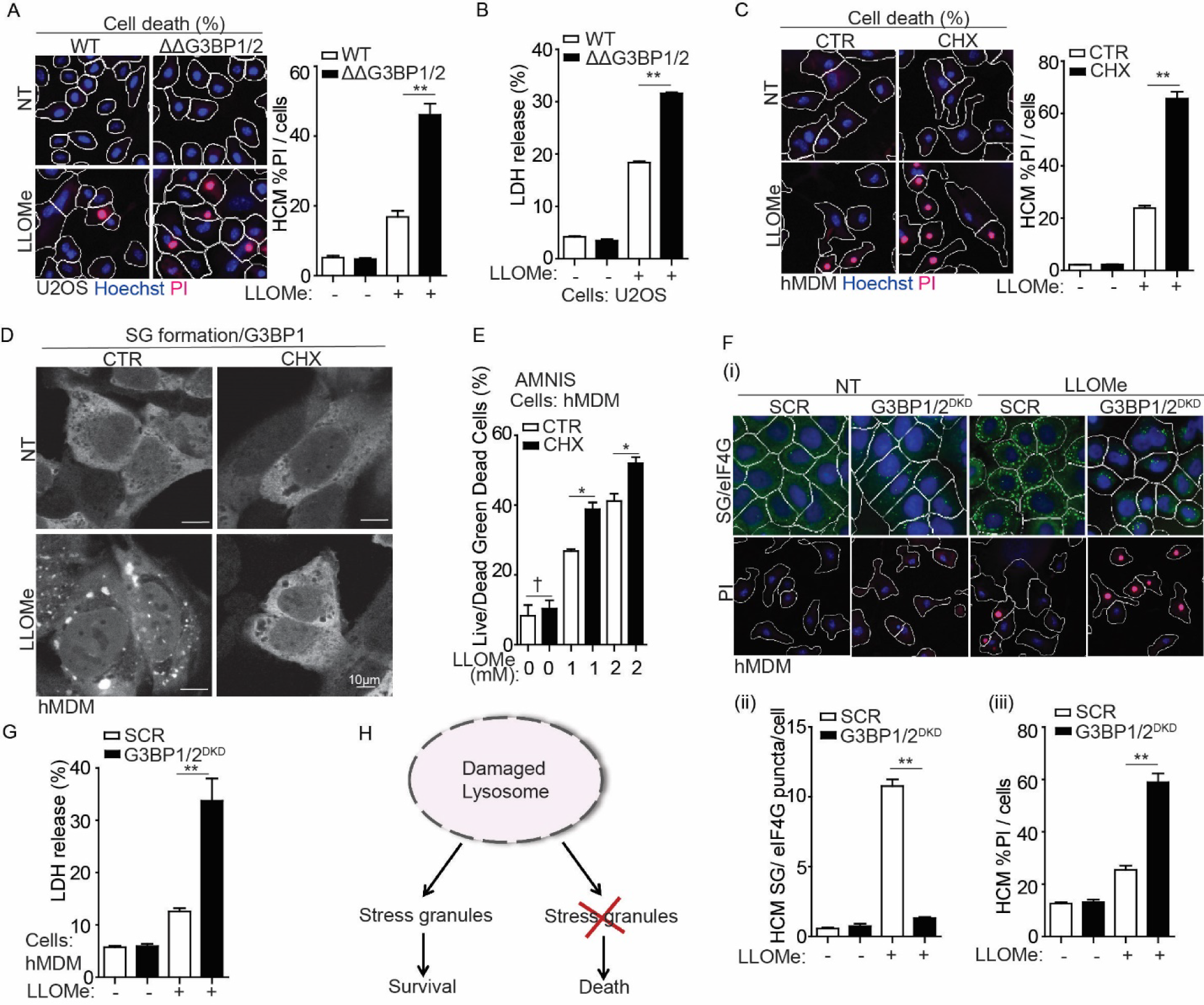
Stress granule formation promotes cell survival in response to lysosomal damage. (A) Quantification by high-content microscopy (HCM) of cell death by a propidium iodide (PI) uptake assay in U2OS wildtype (WT) and G3BP1&2 double knockout (ΔΔG3BP1/2) cells. Cells were treated with 2 mM LLOMe for 30 min, and then stained with propidium iodide (PI) (dead cells) and Hoechst-33342 (total cells). White masks, algorithm-defined cell boundaries (primary objects); red masks, computer-identified PI^+^ nuclei (target objects). (B) Cell death analysis of supernatants of U2OS WT and ΔΔG3BP1/2 cells by a LDH release assay. Cells were treated with 2 mM LLOMe for 30 min. (C) Quantification by HCM of cell death by a PI uptake assay in human peripheral blood monocyte-derived macrophages (hMDM). Cells were treated with 2 mM in the presence or absence of 10 μg/ml cycloheximide (CHX) for 30 min, and then stained with PI (dead cells) and Hoechst-33342 (total cells). (D) Confocal microscopy analysis of G3BP1 (Alexa Fluor 488) in hMDM treated with 2 mM LLOMe with or without CHX for 30 min. Scale bar, 10 μm. (E) Quantification using AMNIS of cell death by Live/Dead^TM^ stain kit in hMDM. Cells were treated with 2 mM LLOMe with or without CHX for 30 min, and then stained using Live/Dead^TM^ stain kit (ThermoFisher). (F) Quantification by HCM of cell death by a PI uptake assay and SG formation by eIF4G in hMDM transfected with scrambled siRNA as control (SCR) or G3BP1 and G3BP2 siRNA for double knockdown (DKD). Cells were treated with 2 mM LLOMe for 30 min, and then stained with PI (dead cells), Hoechst-33342 (total cells) or eIF4G. (i) HCM images: white masks, algorithm-defined cell boundaries; green masks, computer-identified eIF4G puncta; red masks, computer-identified PI^+^ nuclei (target objects); (ii and iii) corresponding HCM quantification. (G) Cell death analysis of supernatants of hMDM transfected with either scrambled siRNA as control (SCR) or G3BP1 and G3BP2 siRNA for double knockdown (DKD) using a LDH release assay. Cells were treated with 2 mM LLOMe for 30 min. (H) Schematic summary of the findings in Figure 1 and S1. CTR, control; NT, untreated cells. Data, means ± SEM (n = 3); HCM: n ≥ 3 (each experiment: 500 valid primary objects/cells per well, ≥5 wells/sample). † p ≥ 0.05 (not significant), *p < 0.05, **p < 0.01, ANOVA. See also Figure S1.

The protective effects of SG formation in response to lysosomal damage were also observed in primary cells using human peripheral blood monocyte-derived macrophages (hMDM). This includes that the significant increase in cell death during LLOMe treatment, as quantified by the PI uptake assay when SG formation was inhibited by cycloheximide in hMDM (Figs. 1C, D). This was further confirmed by measuring the viability of live hMDM (without the fixation) using an AMNIS imaging flow cytometer (Fig. 1E). Knockdown of both G3BP1 and G3BP2 in hMDM (G3BP1/2^DKD^) resulted in a reduction of SG formation as evaluated by a key SG marker, eIF4G puncta, during LLOMe treatment (Figs. 1F (i, ii) and S1H). Elevated cell death as quantified by PI uptake assay (Figs. 1F (i, iii)) and the LDH release assay (Fig. 1G) was detected in G3BP1/2^DKD^ in response to LLOMe treatment. In summary, SG formation is a cytoprotective response to lysosomal damage (Fig. 1H).

### Stress granule formation is controlled by eIF2α pathway but not mTORC1 pathway during lysosomal damage

Considering the significance of SG formation during lysosomal damage, what mechanisms regulate SG formation in response to such damage? SG formation occurs as a consequence of protein translation arrest during cellular stress(Riggs et al., 2020; Youn et al., 2019). eIF2α phosphorylation and mTORC1 inactivation are two key upstream events that lead to protein translation arrest and subsequently trigger SG formation(Cotto et al., 1999; Emara et al., 2012; McCormick et al., 2017). Consistent with our earlier studies(Jia et al., 2018; Jia et al., 2022), we confirmed that LLOMe treatment induced eIF2α phosphorylation and mTORC1 inactivation in a dose-dependent manner in U2OS cells (Fig. S2A). To investigate the role of eIF2α and mTORC1pathways in regulating SG formation upon lysosomal damage, we initially knocked down eIF2α in U2OS cells (eIF2α^KD^) (Fig. 2A). We found that eIF2α is necessary for SG formation upon lysosomal damage, which was reflected by the depletion of SG formation in eIF2α^KD^ cells during LLOMe treatment (Fig. 2A). In addition, we examined mTORC1 activity in eIF2α^KD^ cells by detecting the phosphorylation of its substrate 4EBP1 (Ser65) and found that mTORC1 inactivation was not affected by eIF2α depletion upon lysosomal damage (Fig. 2B). This indicates that eIF2α phosphorylation and mTORC1 inactivation are two uncoupled events during lysosomal damage. This was further confirmed by the lack of change in eIF2α phosphorylation upon lysosomal damage in cells expressing constitutively active RagB^Q99L^, which keeps mTORC1 in an active state(Abu-Remaileh et al., 2017; Sancak et al., 2010) (Fig. 2C). Additionally, SG formation was not affected in cells expressing RagB^Q99L^ in response to lysosomal damage (Fig. 2D). This uncoupled relationship between eIF2α phosphorylation and mTORC1 inactivation in SG formation is also reflected in various cellular stress, including amino acid starvation and arsenic stress. We found that amino acid starvation resulted in mTORC1 inactivation (assessed by mTOR dissociation from the lysosomes(Abu-Remaileh et al., 2017; Jia et al., 2022) but not eIF2α phosphorylation or SG formation in keeping with previous reports(Prentzell et al., 2021; X. Wang et al., 2008) (Figs. S2B, C). In contrast, arsenic stress led to eIF2α phosphorylation and SG formation while activating mTORC1 activity, consistent with earlier studies(Q.Y. Chen et al., 2018; Prentzell et al., 2021; Thedieck et al., 2013) (Figs. S2B, C). The key role of eIF2α phosphorylation in SG formation during lysosomal damage was further demonstrated by the ability of complementing eIF2α WT but not its phosphorylation site mutant (eIF2α S51A)(N.L. Kedersha et al., 1999) in eIF2α^KD^ cells to restore SG formation (Fig. 2E). In summary, eIF2α phosphorylation is a major upstream event for SG formation in response to lysosomal damage (Fig. 2F).

**Figure 2.**
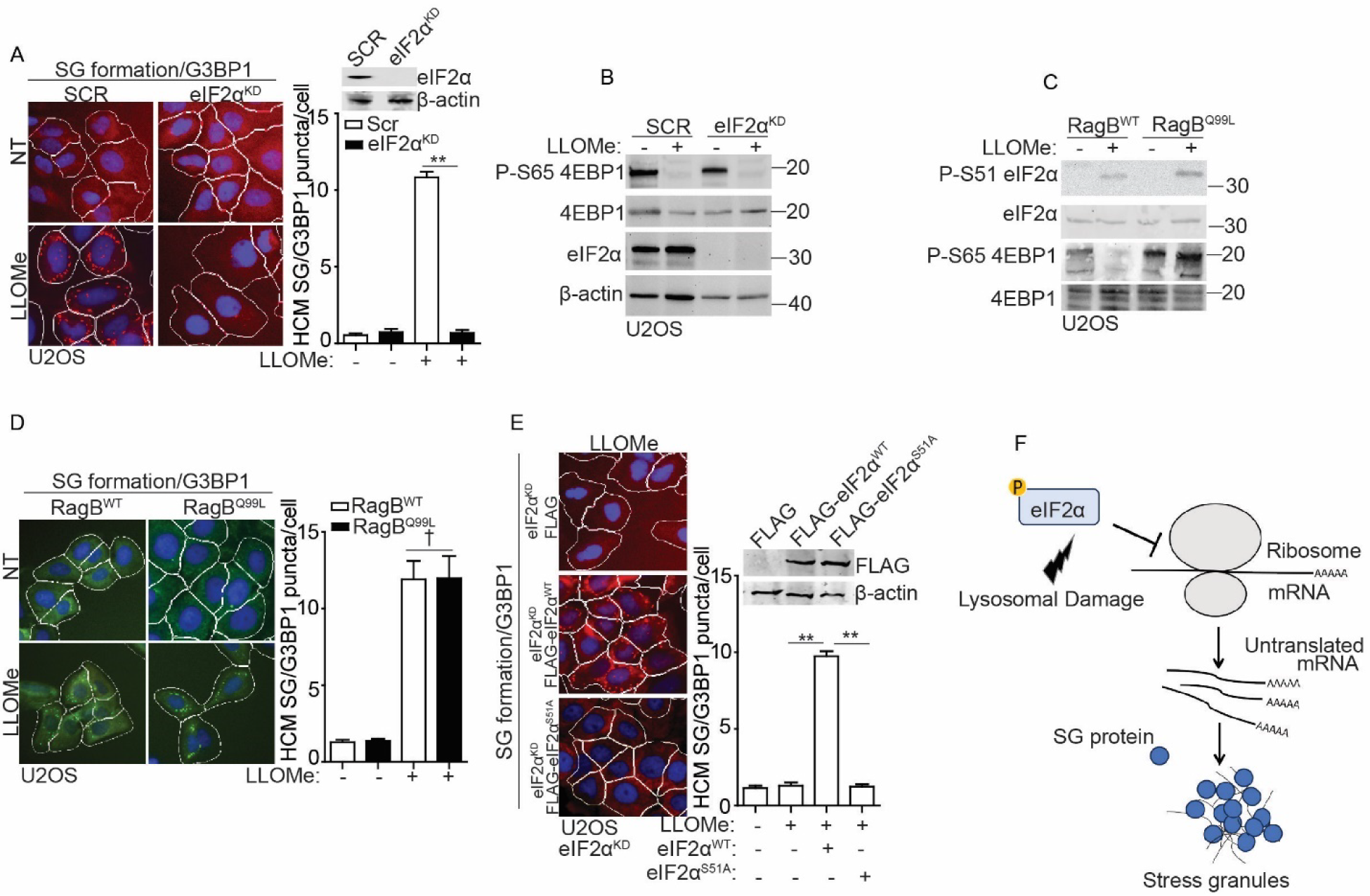
Stress granule formation is controlled by eIF2α pathway but not mTORC1 pathway during lysosomal damage. (A) Quantification by HCM of G3BP1 puncta in U2OS cells transfected with either scrambled siRNA as control (SCR) or eIF2α siRNA for knockdown (eIF2α^KD^). Cells were treated with 2 mM LLOMe for 30 min. White masks, algorithm-defined cell boundaries; red masks, computer-identified G3BP1 puncta. (B) Immunoblot analysis of mTORC1 activity by phosphorylation of 4EBP1 (S65) in U2OS cells transfected with either scrambled siRNA as control (SCR) or eIF2α siRNA for knockdown (eIF2α^KD^). Cells were treated with 2 mM LLOMe for 30 min. (C) Immunoblot analysis of phosphorylation of eIF2α (S51) in U2OS cells overexpressing wildtype RagB (RagB^WT^) or constitutively active RagB mutant (RagB^Q99L^) treated with 2 mM LLOMe for 30 min. (D) Quantification by HCM of G3BP1 puncta in U2OS cells overexpressing wildtype RagB (RagB^WT^) or constitutively active RagB mutant (RagB^Q99L^). Cells were treated with 2 mM LLOMe for 30 min. White masks, algorithm-defined cell boundaries; green masks, computer-identified G3BP1 puncta. (E) Quantification by HCM of G3BP1 puncta in eIF2α knockdown (eIF2α^KD^) U2OS cells transfected with FLAG, FLAG-eIF2α^WT^ or FLAG-eIF2α^S51A^. Cells were treated with 2 Mm LLOMe for 30 min. White masks, algorithm-defined cell boundaries; red masks, computer-identified G3BP1 puncta. (F) Schematic summary of the findings in Figure 2 and S2. NT, untreated cells. Data, means ± SEM (n = 3); HCM: n ≥ 3 (each experiment: 500 valid primary objects/cells per well, ≥5 wells/sample). † p ≥ 0.05 (not significant), **p < 0.01, ANOVA. See also Figure S2.

### Proteomics proximity analysis of eIF2α upon lysosomal damage reveals that its phosphorylation is driven by PKR and PACT

To further investigate the mechanisms that trigger eIF2α phosphorylation in response to lysosomal damage, we conducted a dynamic proteomic analysis using proximity biotinylation. We identified and compared the interacting partners of eIF2α through LC/MS/MS, utilizing APEX2-eIF2α fusion, under both control and lysosomal damage (LLOMe) conditions (for a total of three independent experiments) (Table S1, Tab 1). Volcano plots of this proteomic analysis showed dynamic changes in the proximity of cellular proteins to APEX2-eIF2α during lysosomal damage (Fig. 3A). Within the top twenty candidates showing increased association with eIF2α in response to lysosomal damage, we found the expected candidate PKR (EIF2AK2), which was previously reported by our group as a potential upstream kinase responsible for eIF2α phosphorylation during lysosomal damage(Jia et al., 2022)(Fig. 3A). Interestingly, the activator of PKR, PACT (PRKRA)(Patel et al., 1998), prominently emerged with the most significant fold increase following lysosomal damage (Fig. 3A). PACT is known to facilitate the stress-induced phosphorylation and activation of PKR through direct interaction(Patel et al., 1998; Singh et al., 2012). This interaction disrupts PKR’s self-inhibition, leading to PKR autophosphorylation including at Thr446, which converts it into its fully active form capable of phosphorylating protein substrates, such as eIF2α(Chukwurah et al., 2021; Sadler et al., 2007). We confirmed increased interactions of PKR and PACT with eIF2α upon lysosomal damage by co-immunoprecipitation (co-IP) between FLAG-eIF2α and endogenous PKR and PACT (Fig. 3B). Next, we examined whether PKR and PACT are functionally necessary for eIF2α phosphorylation triggered by lysosomal damage. Previously, we knocked down four widely recognized upstream kinases of eIF2α (HRI, PKR, PERK and GCN2)(Pakos-Zebrucka et al., 2016), and found that only the knockdown of PKR resulted in the inhibition of eIF2α phosphorylation and SG formation(Jia et al., 2022). To confirm these findings, we generated a CRISPR knockout of PKR (PKR^KO^) in SG reporter cells (U2OS G3BP1-GFP). In these PKR^KO^ cells, the formation of SG induced by lysosomal damage was completely inhibited, as quantified by the puncta of G3BP1-GFP using HCM (Fig. 3C). In line with this, the phosphorylation of eIF2α and PKR was also abolished (Fig. 3D). Conversely, the overexpression of PKR in PKR^KO^ cells led to a restoration of phosphorylation of eIF2α and PKR during lysosomal damage (Fig. 3D). Recently, MARK2 was identified as the fifth kinase responsible for eIF2α phosphorylation in response to proteotoxic stress(Lu et al., 2021). However, we found that MARK2 did not regulate eIF2α phosphorylation during lysosomal damage (Fig. S3A). Next, we examined PKR’s activator, PACT by knocking down PACT in U2OS cells (PACT^KD^). We observed a decrease in PKR activation, eIF2α phosphorylation and SG formation observed in PACT^KD^ cells during lysosomal damage (Fig. 3E). This finding aligns with the role of PKR in controlling eIF2α phosphorylation and SG formation. Thus, both PKR and its activator PACT regulate eIF2α phosphorylation for SG formation in response to lysosomal damage.

**Figure 3.**
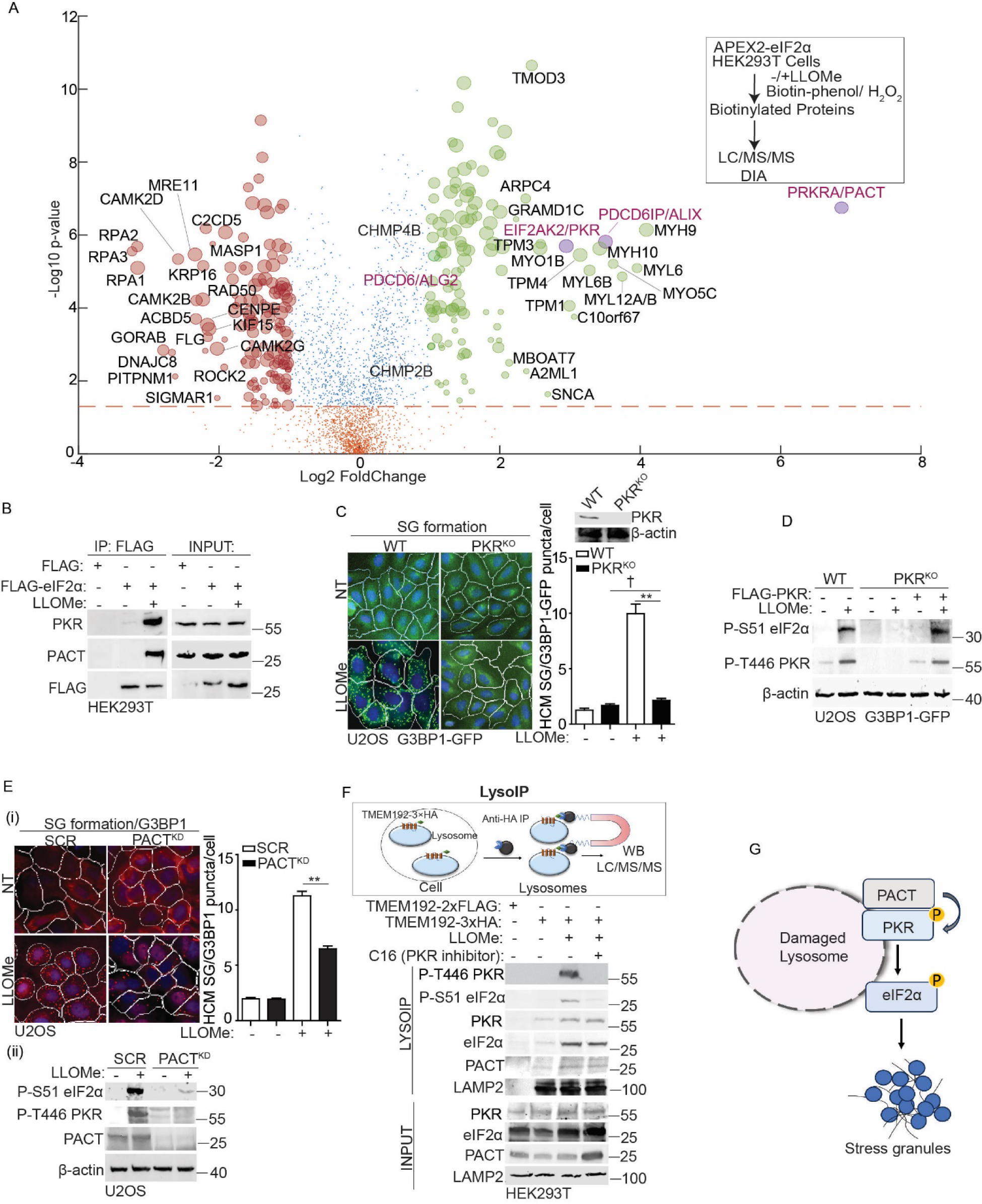
PKR and its activator PACT regulate eIF2α phosphorylation on damaged lysosomes. (A) Quantitative liquid chromatography-tandem mass spectrometry (LC/MS/MS) using the data-independent acquisition (DIA) technique to identify eIF2α binding partners that were proximity-biotinylated by APEX2-eIF2α during lysosomal damage (1mM LLOMe for 1h). Scatter (volcano) plot shows log2 fold change (LLOMe/CTR; spectral counts) and –log10 p value for the proteins identified and quantified in three independent experiments. Green dots indicate increase in proximity to eIF2α (log2 fold change ≥ 1), and red dots indicate decrease in proximity to eIF2α (log2 fold change ≤ −1) during LLOMe treatment. Orange dots indicate values below the statistical significance cut-off (P ≥ 0.05). Bubble size represents a normalized value for the total amount of spectral counts for the protein indicated. PACT, PKR and ALIX proteins are highlighted as purple circles (see Table S1). (B) Co-IP analysis of interactions between eIF2α and PKR/PACT during lysosomal damage. HEK293T cells expressing FLAG (control) or FLAG-eIF2α were treated with 1 mM LLOMe for 30 min. Cell lysates were immunoprecipitated with anti-FLAG antibody and immunoblotted for indicated proteins. (C) Quantification by HCM of G3BP1-GFP puncta in wildtype (WT) or PKR knockout (PKR^KO^) U2OS G3BP1-GFP cells. Cells were treated with 2 mM LLOMe for 30 min. White masks, algorithm-defined cell boundaries; green masks, computer-identified G3BP1 puncta. (D) Immunoblot analysis of phosphorylation of eIF2α (S51) and PKR (T446) in WT or PKR^KO^ U2OS G3BP1-GFP cells, as well as in cells overexpressing FLAG-PKR in PKR^KO^ U2OS G3BP1-GFP cells. Cells were treated with 2 mM LLOMe for 30 min. (E) (i) Quantification by HCM of G3BP1 puncta in U2OS cells transfected with either scrambled siRNA as control (SCR) or PACT siRNA for knockdown (PACT^KD^). Cells were treated with 2 mM LLOMe for 30 min. White masks, algorithm-defined cell boundaries; red masks, computer-identified G3BP1 puncta; (ii) Immunoblot analysis of phosphorylation of eIF2α (S51) and PKR (T446) in SCR or PACT^KD^ cells; 2 mM LLOMe for 30 min. (F) Analysis of proteins associated with purified lysosomes (LysoIP; TMEM192-3xHA) from HEK293T cells treated with 1 mM LLOMe in the presence or absence of 210 nM imidazolo-oxindole C16 for 1h. TMEM192-2xFLAG, control. (G) Schematic summary of the findings in Figure 3 and S3. NT, untreated cells. Data, means ± SEM (n = 3); HCM: n ≥ 3 (each experiment: 500 valid primary objects/cells per well, ≥5 wells/sample). † p ≥ 0.05 (not significant), **p < 0.01, ANOVA. See also Figure S3.

### PKR and PACT control eIF2α phosphorylation on damaged lysosomes

We previously performed proteomic analyses of lysosomes that were purified using LysoIP(Jia et al., 2022), a well-established approach to isolate lysosomes by the lysosomal membrane protein TMEM192(Abu-Remaileh et al., 2017; Jia et al., 2020c). These analyses indicate the presence of PKR, PACT and eIF2α on lysosomes (Fig. S3B). This finding is further supported by similar results from LysoIP proteomic analysis conducted by other research groups(Eapen et al., 2021; Wyant et al., 2018)(Fig. S3B). Using LysoIP immunoblotting, we confirmed the presence of PKR, PACT and eIF2α on lysosomes and found an elevation in their association with damaged lysosomes (Fig. 3F). We also observed that the phosphorylation of both PKR and eIF2α occurred on damaged lysosomes. Notably, this effect was effectively blocked by a specific PKR’s inhibitor, imidazolo-oxindole C16, known for its ability to inhibit PKR’s autophosphorylation by binding to PKR’s ATP-binding pocket(Gal-Ben-Ari et al., 2019; Jammi et al., 2003; Tronel et al., 2014) (Fig. 3F). Moreover, through confocal fluorescence microscopy, we detected an increased association of PKR, PACT and eIF2α with damaged lysosomes (Figs. S3C-E). In summary, we conclude that PKR and its activator PACT regulate eIF2α phosphorylation on damaged lysosomes (Fig. 3G).

### ALIX and ALG2 are required for stress granule formation by sensing calcium release from damaged lysosomes

In our proteomic analysis of eIF2α binding partners (Fig. 3A), we observed an increased association between eIF2α and ESCRT components such as ALIX, CHMP2B and CHMP4B following lysosomal damage. Specially, ALIX showed a more than10 fold increase (Fig. 3A). We next determined whether these ESCRT components were involved in eIF2α phosphorylation and SG formation triggered by lysosomal damage. Upon lysosomal damage, we observed a significant reduction in SG formation upon knockdown of ALIX in U2OS cells (ALIX^KD^), as quantified by G3BP1 puncta using HCM (Figs. 4A, C). This was also reflected in the decreased phosphorylation of eIF2α and PKR in ALIX^KD^ cells during LLOMe treatment (Figs. 4B, D), indicating an impact of ALIX on the upstream signaling of SG formation. However, the knockdown of CHM2B or CHMP4B had no discernible effect on SG formation and its upstream events (Figs. S4A, B). Previous studies showed that the depletion of both ALIX and TSG101 effectively impedes lysosomal repair by eliminating ESCRT recruitment(Niekamp et al., 2022; Radulovic et al., 2018; Skowyra et al., 2018). We found that TSG101 has no effect on the regulation of SG formation upon lysosomal damage. This is supported by the absence of any significant changes in SG formation and eIF2α phosphorylation in TSG101 knockdown U2OS cells (TSG101^KD^) (Figs. 4C, D). ALIX has been reported to sense lysosomal damage through the detection of calcium leakage, which is facilitated by its calcium binding partner, ALG2(W. Chen et al., 2024; Jia et al., 2020a; Niekamp et al., 2022; Skowyra et al., 2018). Notably, ALG2 exhibited increased proximity to eIF2α upon lysosomal damage (Fig. 3A). To further determine the regulatory role of ALIX in SG formation upon lysosomal damage, we utilized BAPTA-AM, the calcium chelator and ALG2 knockdown U2OS cells (ALG2^KD^) to prevent the recruitment of ALIX to damaged lysosomes as previously reported(Jia et al., 2020a; Skowyra et al., 2018). This was confirmed by the observed decrease in ALIX puncta formation upon lysosomal damage in cells treated with BAPTA-AM or in ALG2^KD^ cells (Fig. S4C). Importantly, we also observed a significant reduction in SG formation and eIF2α phosphorylation in cells treated with BAPTA-AM, or in ALG2^KD^ cells during lysosomal damage (Figs. 4E, F). Thus, we conclude that ALIX and its partner ALG2 modulate eIF2α phosphorylation by sensing calcium leakage as lysosomal damage signal, thereby initiating SG formation (Fig. 4G).

**Figure 4.**
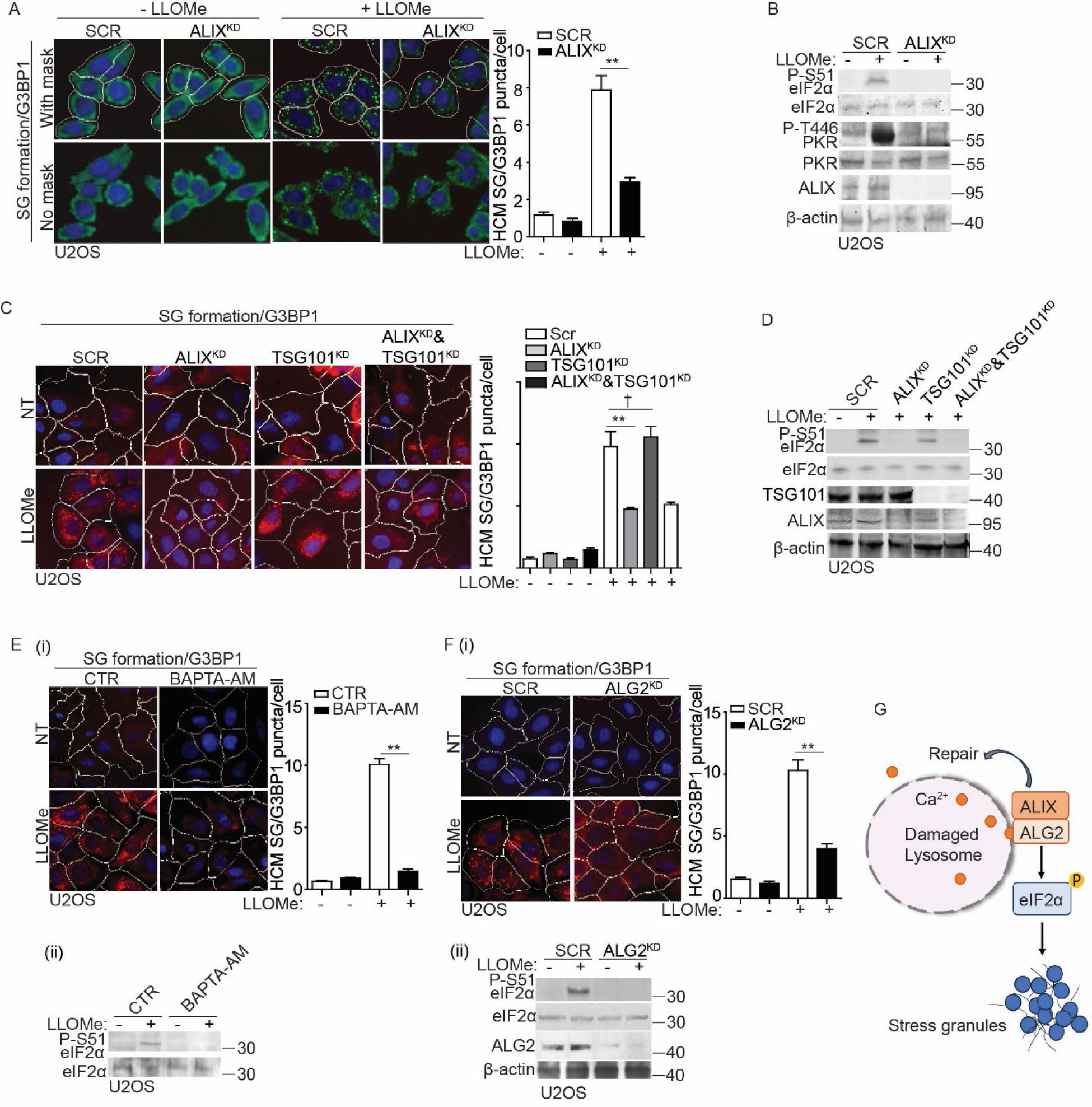
ALIX and ALG2 are required for stress granule formation by sensing calcium release from damaged lysosomes. (A) Quantification by HCM of G3BP1 puncta in U2OS cells transfected with either scrambled siRNA as control (SCR) or ALIX siRNA for knockdown (ALIX^KD^). Cells were treated with 2 mM LLOMe for 30 min. White masks, algorithm-defined cell boundaries; green masks, computer-identified G3BP1 puncta. (B) Immunoblot analysis of phosphorylation of eIF2α (S51) and PKR (T446) in U2OS cells transfected with either scrambled siRNA as control (SCR) or ALIX siRNA for knockdown (ALIX^KD^). Cells were treated with 2 mM LLOMe for 30 min. (C) Quantification by HCM of G3BP1 puncta in U2OS cells transfected with scrambled siRNA as control (SCR), ALIX siRNA for knockdown (ALIX^KD^) or TSG101 siRNA for knockdown (TSG101^KD^). Cells were treated with 2 mM LLOMe for 30 min. White masks, algorithm-defined cell boundaries; red masks, computer-identified G3BP1 puncta. (D) Immunoblot analysis of phosphorylation of eIF2α (S51) in U2OS cells transfected with scrambled siRNA as control (SCR), ALIX siRNA for knockdown (ALIX^KD^) or TSG101 siRNA for knockdown (TSG101^KD^). Cells were treated with 2 mM LLOMe for 30 min. (E) (i) Quantification by HCM of G3BP1 puncta in U2OS cells pre-treated with 15 µM BAPTA-AM for 1 h, subjected to 2 mM LLOMe treatment for 30 min. White masks, algorithm-defined cell boundaries; red masks, computer-identified G3BP1 puncta. (ii) Immunoblot analysis of phosphorylation of eIF2α (S51) in U2OS cells as described in (i). (F) (i) Quantification by HCM of G3BP1 puncta in U2OS cells transfected with scrambled siRNA as control (SCR), or ALG2 siRNA for knockdown (ALG2^KD^). Cells were treated with 2 mM LLOMe for 30 min. White masks, algorithm-defined cell boundaries; red masks, computer-identified G3BP1 puncta. (ii) Immunoblot analysis of phosphorylation of eIF2α (S51) in U2OS cells as described in (i). (G) Schematic summary of the findings in Figure 4 and S4. NT, untreated cells. CTR, control. Data, means ± SEM (n = 3); HCM: n ≥ 3 (each experiment: 500 valid primary objects/cells per well, ≥5 wells/sample). † p ≥ 0.05 (not significant), **p < 0.01, ANOVA. See also Figure S4.

### ALIX associates with PKR and PACT in response to lysosomal damage

Given that eIF2α phosphorylation is initiated by its upstream kinase PKR, and its activator PACT (Fig. 3), our subsequent investigation delved into exploring the relationship among ALIX, PKR and PACT. Using the Co-IP assay, we tested the interaction between FLAG-ALIX and endogenous PKR and PACT. Their interactions were notably enhanced following treatment with LLOMe (Fig. 5A). ALIX is composed of three distinct domains: Bro1 domain, V domain, and proline-rich domain (PRD) (Fig. 5B). These domains have the potential to remain inactive due to intramolecular interactions but can be activated through interaction with ALG2 in a calcium-dependent manner(Maki et al., 2016; Scheffer et al., 2014; Sun et al., 2015; Vietri et al., 2020) (Fig. 5B). Next, we generated the domain deletions of ALIX (Fig. 5B(i)). The mapping analysis of ALIX domains necessary for binding to PKR and PACT revealed the indispensable role of the V domain in their interaction (Fig. 5C). Additionally, increased associations among full-length ALIX, PKR and PACT were observed upon LLOMe treatment (Fig. 5C), suggesting that lysosomal damage activates ALIX by releasing its V domain for association with PKR and PACT. This is corroborated by the interaction of the V domain of ALIX with PKR and PACT, even in cells that were not subjected to lysosome damage induced by LLOMe (Fig. 5C). The interactions between ALIX and PKR, as well as ALIX and PACT, were also predicted using the protein-protein docking server HDOCK, resulting in scores of 0.8398 for ALIX-PKR and 0.8325 for ALIX-PACT, reflecting a high level of confidence (Yan et al., 2020) (Figs. S5A, B). Furthermore, through confocal fluorescence microscopy, we observed the association among ALIX, PKR and PACT during lysosomal damage (Fig. S5C). Thus, ALIX interacts with PKR and PACT in response to lysosomal damage.

**Figure 5.**
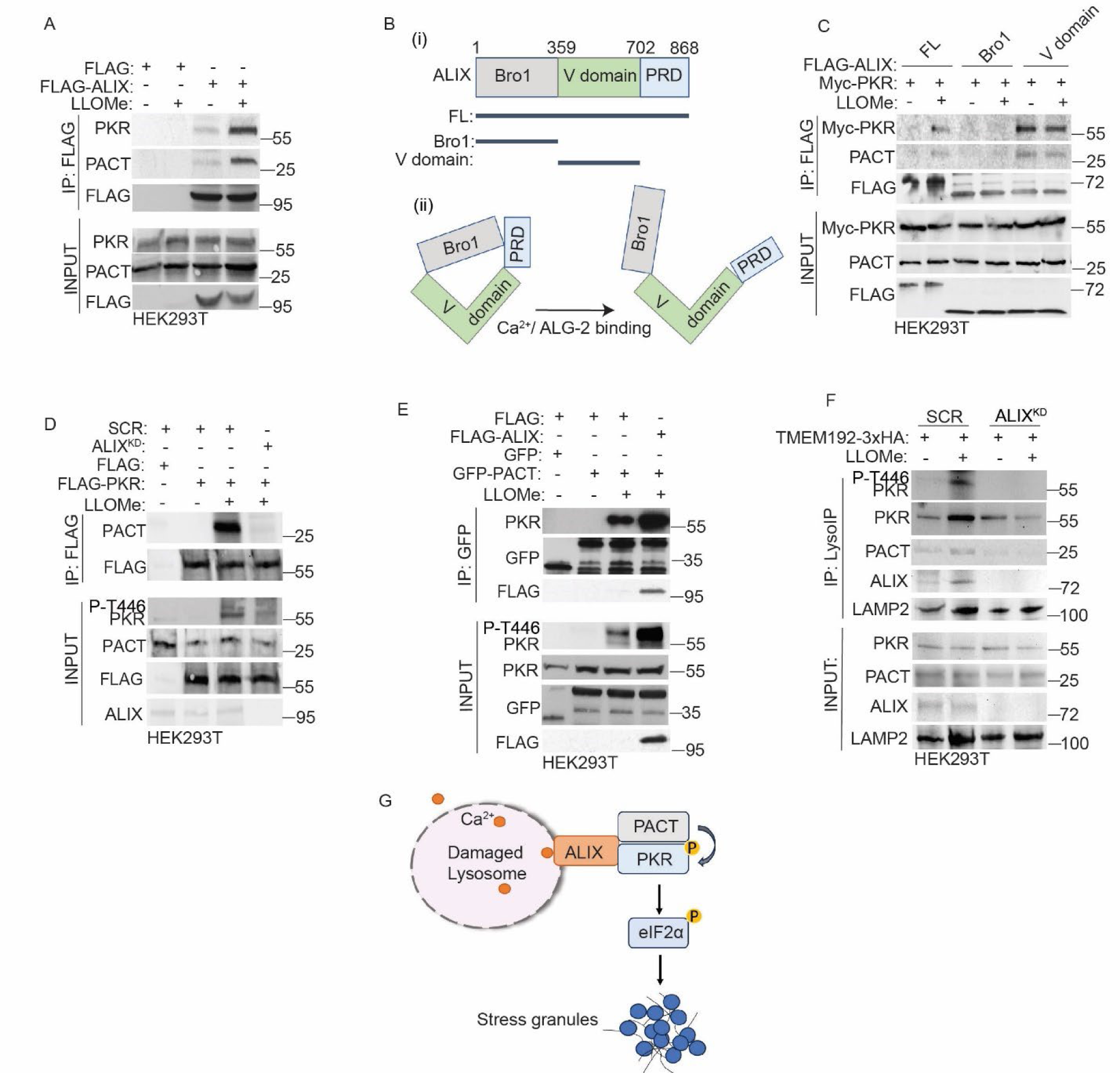
ALIX promotes the association between PKR and its activator PACT on damaged lysosomes. (A) Co-IP analysis of interactions among ALIX, PKR and PACT during lysosomal damage. HEK293T cells expressing FLAG (control) or FLAG-ALIX were treated with 1 mM LLOMe for 30 min. Cell lysates were immunoprecipitated with anti-FLAG antibody and immunoblotted for indicated proteins. (B) (i) Schematic diagram of ALIX mutants used in this study. FL (full length); Bro1 (Bro1 domain); V domain; PRD (proline-rich domain). Numbers, residue positions. (ii) Schematic illustration of the Ca^2+^/ALG-2-induced open conformation of ALIX. (C) Co-IP analysis of interactions among ALIX mutants, PKR and PACT during lysosomal damage. HEK293T cells expressing FLAG tagged ALIX mutants and Myc-PKR were treated with 1 mM LLOMe for 30 min. Cell lysates were immunoprecipitated with anti-FLAG antibody and immunoblotted for indicated proteins. (D) Co-IP analysis of interactions between FLAG-PKR and PACT in HEK293T cells transfected with scrambled siRNA as control (SCR), or ALIX siRNA for knockdown (ALIX^KD^) during lysosomal damage. Cells were treated with 1 mM LLOMe for 30 min. Cell lysates were immunoprecipitated with anti-FLAG antibody and immunoblotted for indicated proteins. (E) Co-IP analysis of interactions between PKR and GFP-PACT in HEK293T cells transfected with FLAG, or FLAG-ALIX during lysosomal damage. Cells were treated with 1 mM LLOMe for 30 min. Cell lysates were immunoprecipitated with anti-GFP antibody and immunoblotted for indicated proteins. (F) Analysis of proteins associated with purified lysosomes (LysoIP; TMEM192-3xHA) from HEK293T cells transfected with scrambled siRNA as control (SCR), or ALIX siRNA for knockdown (ALIX^KD^). Cells were treated with 1 mM LLOMe for 30 min. (G) Schematic summary of the findings in Figure 5 and S5. One of three independent Western Blot experiments shown. See also Figure S5.

### ALIX promotes the association between PKR and its activator PACT on damaged lysosomes

Next, we quantified by HCM the ALIX puncta response to lysosomal damage in cells where PKR or PACT had been knocked down. We observed that the presence or absence of PKR and PACT did not affect ALIX response to lysosomal damage (Fig. S5D). This suggests that ALIX may potentially precede PKR and PACT for eIF2α phosphorylation upon lysosomal damage. Considering the decrease in the phosphorylation of PKR in ALIX^KD^ cells and the increased association among ALIX, PKR and PACT following lysosomal damage (Figs. 4B, D, 5A), we hypothesize that ALIX regulates PKR phosphorylation by modulating the association between PKR and its activator PACT during lysosomal damage. Using co-IP assays, we confirmed the formation of complexes between FLAG-PKR and endogenous PACT during lysosomal damage (Fig. 5D). However, this interaction was reduced in ALIX^KD^ HEK293T cells (Fig. 5D), resulting in decreased PKR phosphorylation during LLOMe treatment. Conversely, the overexpression of ALIX led to a further enhancement in the increased association between GFP-PACT and endogenous PKR, and this was accompanied by an increase in PKR phosphorylation during lysosomal damage (Fig. 5E). These data indicates that ALIX is essential for PKR phosphorylation by controlling the interaction between PKR and PACT during lysosomal damage. Next, we examined whether this regulatory event occurred on damaged lysosomes by conducting LysoIP immunoblotting in ALIX^KD^ HEK293T cells. In this assay, we observed that ALIX^KD^ HEK293T cells no longer displayed PKR phosphorylation on damaged lysosomes, accompanied by a reduced recruitment of PKR and PACT to lysosomes, as determined by Western blot analysis of lysosomes isolated using LysoIP (Fig. 5F). This suggests that ALIX is responsible for the recruitment and regulation of PKR and PACT on damaged lysosomes. In summary, we conclude that ALIX recruits PKR and its activator PACT to damaged lysosomes and regulates the activation of PKR by enhancing its association with PACT, consequently leading to eIF2α phosphorylation and SG formation (Fig. 5G).

### Galectin-3 inhibits stress granule formation by reducing the association between PKR and PACT during lysosomal damage

Previously, we reported that galectin-3 (Gal3), a β-galactoside-binding protein that recognizes damage-exposed glycan, can recruit ALIX to damaged lysosomes and promote ESCRT function for lysosomal repair and restoration(Jia et al., 2020a). We examined whether Gal3 is involved in the regulatory process of SG formation during lysosomal damage. In U2OS cells subjected to Gal3 knockdown (Gal3^KD^), we observed an elevated level of SG formation, quantified by the formation of G3BP1 puncta using HCM (Fig. 6A). This result was consistent with our earlier reported increase in SGs in Gal3 knockout HeLa cells(Jia et al., 2022). We further detected the upstream signaling events leading to SG formation in Gal3^KD^ U2OS cells and observed an increase in the phosphorylation of PKR and eIF2α in the absence of Gal3 following LLOMe treatment (Fig. 6B). These data indicate that Gal3 has a negative effect on the activation of PKR and eIF2α, thereby affecting SG formation during lysosomal damage. We next tested the relationship among Gal3, PKR and PACT. The Co-IP results showed that Gal3 can be in protein complexes with ALIX, PKR and PACT upon lysosomal damage (Fig. 6C). When we investigated whether Gal3 can control the association between PKR and PACT, we found an increase in their association in the absence of Gal3 (Fig. 6D). This was further confirmed by the increased PKR phosphorylation under the same conditions. On the contrary, when Gal3 was overexpressed, it led to a reduction in the interaction between PKR and PACT, consequently resulting in reduced PKR phosphorylation upon LLOMe treatment (Fig. 6E). We interpret the inhibitory role of Gal3 in the association between PKR and PACT as a result of their competition for ALIX. In keeping with this interpretation, we observed a reduced interaction among ALIX, PACT and PKR in Gal3-overexpressing cells during LLOMe treatment (Fig. 6F). However, when we overexpressed the Gal3^R186S^ mutant, which has been previously shown to lose the ability to recognize damaged lysosomes(Aits et al., 2015; Jia et al., 2020a), it failed to regulate the protein complex of ALIX, PACT and PKR upon lysosomal damage (Fig. 6F). Moreover, given our previous finding that Gal3 facilitates ESCRT-mediated lysosomal repair via ALIX(Jia et al., 2020a), these observations provide evidence of Gal3’s role in balancing ALIX-mediated lysosomal repair and ALIX-mediated SG formation (Fig. 6G). Thus, we conclude that the recruitment of Gal3 to damaged lysosomes plays an inhibitory effect on the regulation of the upstream processes of SG formation by decreasing the association between PKR and PACT (Fig. 6G).

**Figure 6.**
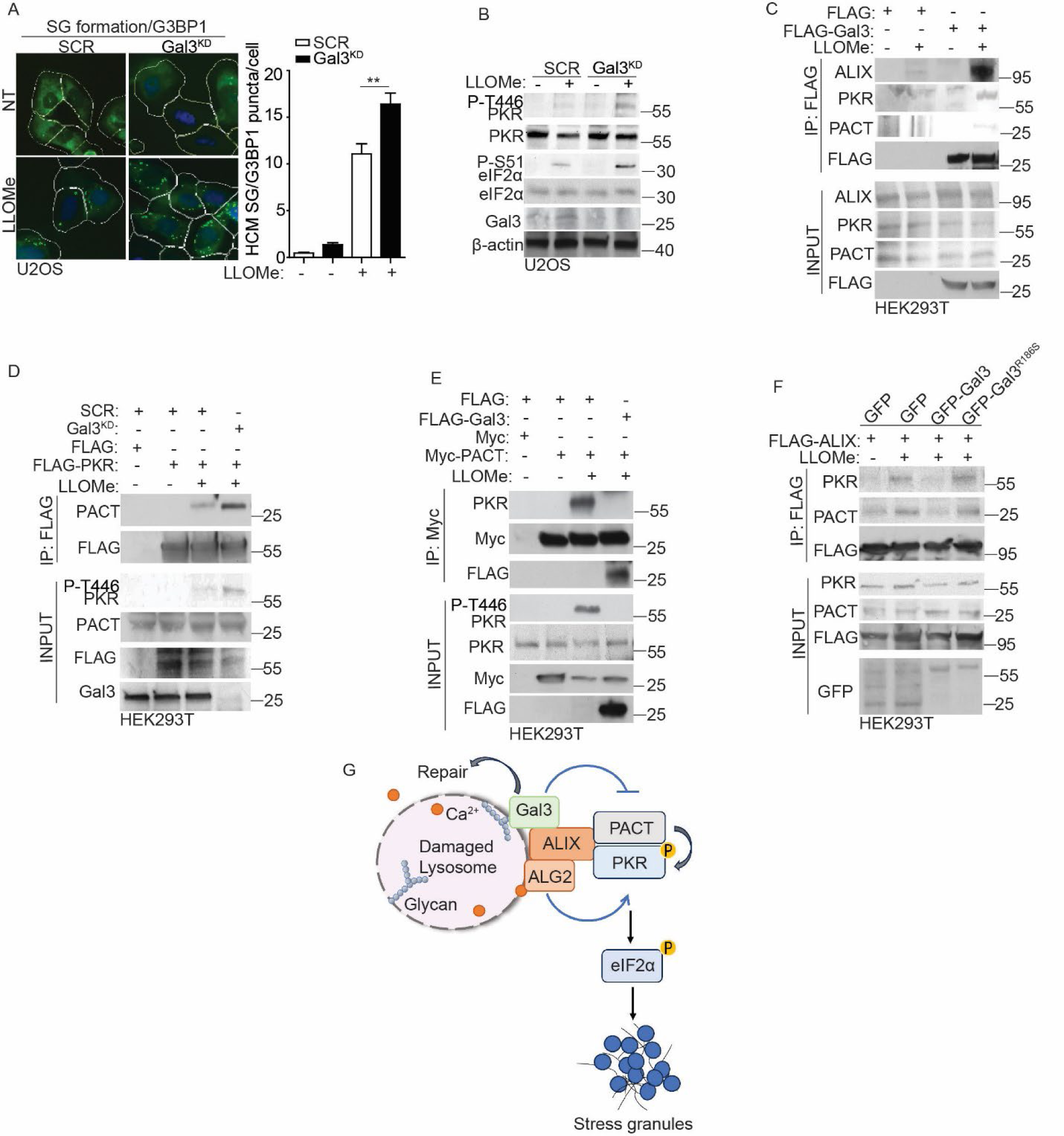
Galectin-3 inhibits stress granule formation by reducing the association between PKR and PACT during lysosomal damage. (A) Quantification by HCM of G3BP1 puncta in U2OS cells transfected with scrambled siRNA as control (SCR), or galectin-3 (Gal3) siRNA for knockdown (Gal3^KD^). Cells were treated with 2 mM LLOMe for 30 min. White masks, algorithm-defined cell boundaries; green masks, computer-identified G3BP1 puncta. (B) Immunoblot analysis of phosphorylation of eIF2α (S51) and PKR (T446) in U2OS cells transfected with scrambled siRNA as control (SCR), or galectin-3 (Gal3) siRNA for knockdown (Gal3^KD^), subjected to 2 mM LLOMe treatment for 30 min. (C) Co-IP analysis of interactions among FLAG-Gal3, ALIX, PKR and PACT in HEK293T cells during lysosomal damage. Cells were treated with 1 mM LLOMe for 30 min. Cell lysates were immunoprecipitated with anti-FLAG antibody and immunoblotted for indicated proteins. (D) Co-IP analysis of interactions between FLAG-PKR and PACT in HEK293T cells transfected with scrambled siRNA as control (SCR), or Gal3 siRNA for knockdown (Gal3^KD^) during lysosomal damage. Cells were treated with 1 mM LLOMe for 30 min. Cell lysates were immunoprecipitated with anti-FLAG antibody and immunoblotted for indicated proteins. (E) Co-IP analysis of interactions between Myc-PACT and PKR in HEK293T cells transfected with FLAG, or FLAG-Gal3 during lysosomal damage. Cells were treated with 1 mM LLOMe for 30 min. Cell lysates were immunoprecipitated with anti-Myc antibody and immunoblotted for indicated proteins. (F) Co-IP analysis of interactions among FLAG-ALIX, PKR and PACT in HEK293T cells transfected with GFP, GFP-Gal3 or GFP-Gal3^R186S^ during lysosomal damage. Cells were treated with 1 mM LLOMe for 30 min. Cell lysates were immunoprecipitated with anti-FLAG antibody and immunoblotted for indicated proteins. (G) Schematic summary of the findings in Figure 6. NT, untreated cells. Data, means ± SEM (n = 3); HCM: n ≥ 3 (each experiment: 500 valid primary objects/cells per well, ≥5 wells/sample). **p < 0.01, ANOVA. One of three independent Western Blot experiments shown.

### Stress granule formation promotes cell survival in response to lysosomal damage during disease states

Lysosomal damage serves as both a cause and consequence of many disease conditions, including infectious and neurodegenerative diseases(Amaral et al., 2023; Ballabio et al., 2020; Bonam et al., 2019; Fehrenbacher et al., 2005). We tested whether the above molecular and cellular processes that transduce lysosomal damage signals to induce SG formation are important for cell survival in disease contexts. Lysosomal damage can occur from viral infections including those caused by non-enveloped adenovirus and enveloped SARS-CoV-2 infections (Aits et al., 2013; Barlan et al., 2011; Daussy et al., 2020; Thurston et al., 2012; Fengjuan Wang et al., 2018). Adenovirus enters cells through endocytosis and damages lysosomes by releasing its protease, enabling access to the cytosol for replication(A. Barlan et al., 2011; Greber et al., 1996; Pied et al., 2022; Wiethoff et al., 2015). We employed the wildtype human adenovirus species C2 (HAdV-C2^WT^) and its protease-deficient mutant TS1 (HAdV-C2^TS1^), the latter lacking the ability to damage lysosomes(Gallardo et al., 2021; Greber et al., 1996; Martinez et al., 2015). U2OS cells were infected with either HAdV-C2^WT^ or HAdV-C2^TS1^ for 1h and lysosomal damage marker LysoTracker Red (LTR), which measures lysosomal acidification(Chazotte, 2011; Jia et al., 2020a; Pierzyńska-Mach et al., 2014) was quantified by HCM in these infected cells. Consistent with earlier findings(Luisoni et al., 2015; Martinez et al., 2015; Pied et al., 2022), HAdV-C2^WT^ led to a reduction in LTR^+^ profiles, whereas HAdV-C2^TS1^ did not show a similar effect (Fig. S6A). Additionally, SG formation and eIF2α phosphorylation were detected in cells infected with HAdV-C2^WT^ but not in those infected with HAdV-C2^TS1^ (Figs. 7A, B). These results imply that lysosomal damage triggered by HAdV-C2 infection can activate the eIF2α pathway, resulting in SG formation. We tested whether SG formation is important for cell survival during HAdV-C2 infection. In SG-deficient U2OS (ΔΔG3BP1/2) cells compared to wildtype U2OS cells, we observed an elevated level of cell death by PI uptake assay during HAdV-C2^WT^ infection (Fig. 7C). Moreover, our recent publication(Jia et al., 2022) showed that lysosomal damage induced by the expression of SARS-CoV-2 ORF3a protein can also trigger SG formation. Following the overexpression of SARS-CoV-2 ORF3a in U2OS cells, a notable rise in cell death was observed through LDH release assay in ΔΔG3BP1/2 cells compared to control cells (Fig. 7D). Thus, SG formation triggered by lysosomal damage emerges as a crucial process for cell survival during viral infection.

**Figure 7.**
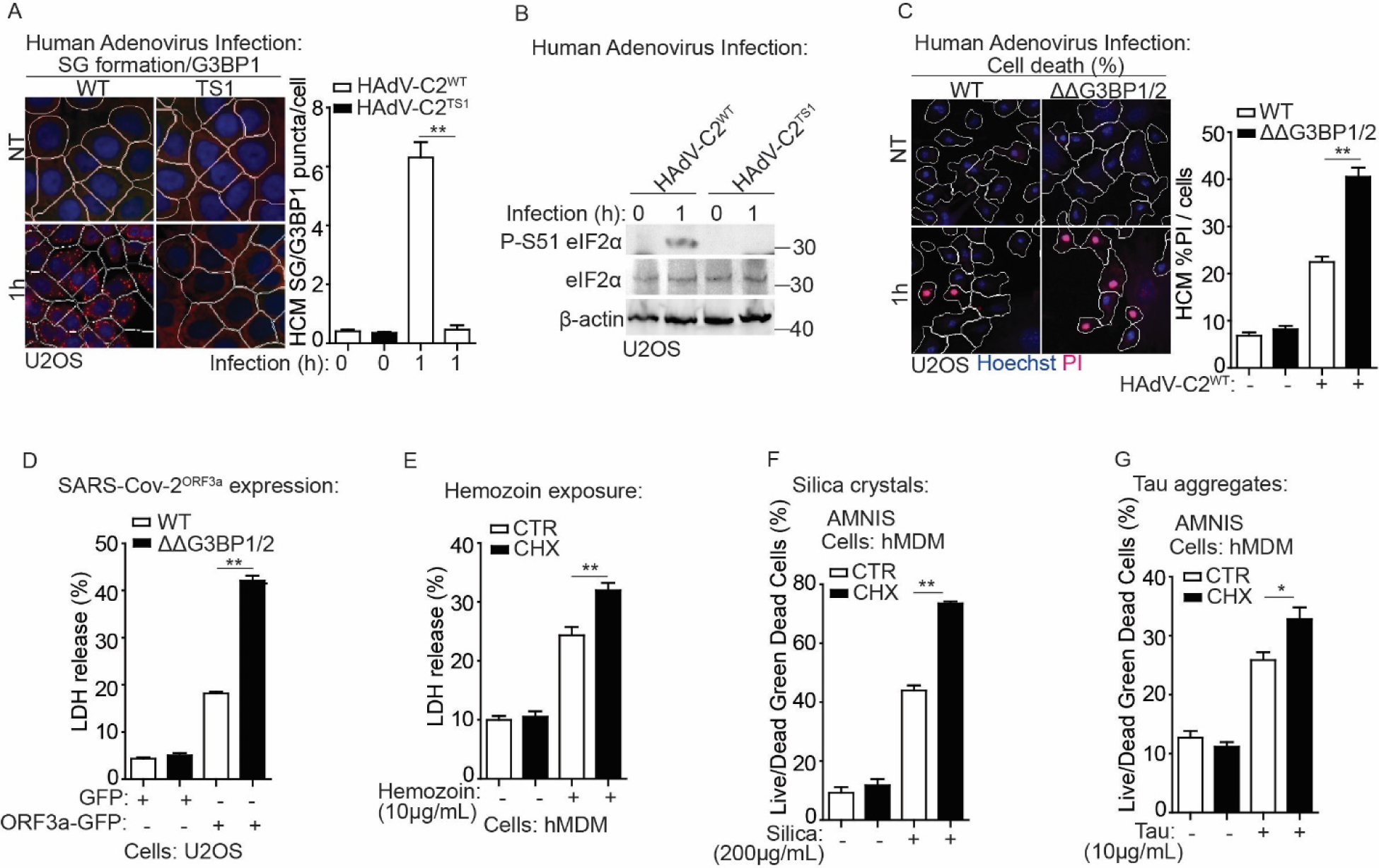
Stress granule formation promotes cell survival in response to lysosomal damage during disease states. (A) Quantification by HCM of G3BP1 puncta in U2OS cells infected with wildtype human adenovirus C2 (HAdV-C2^WT^) or C2 TS1 mutant (HAdV-C2^TS1^) at MOI=10 for 1h. White masks, algorithm-defined cell boundaries; red masks, computer-identified G3BP1 puncta. (B) Immunoblot analysis of phosphorylation of eIF2α (S51) in U2OS cells infected with wildtype human adenovirus C2 (HAdV-C2^WT^) or C2 TS1 mutant (HAdV-C2^TS1^) at MOI=10 for 1h. (C) Quantification by HCM of cell death by a propidium iodide (PI) uptake assay in U2OS wildtype (WT) and G3BP1&2 double knockout (ΔΔG3BP1/2) cells during adenovirus infection. Cells were infected with wildtype human adenovirus C2 (HAdV-C2^WT^) at MOI=10 for 1h, and then stained with propidium iodide PI (dead cells) and Hoechst-33342 (total cells). White masks, algorithm-defined cell boundaries; red masks, computer-identified PI^+^ nuclei. (D) Cell death analysis of supernatants of U2OS WT and ΔΔG3BP1/2 cells by a LDH release assay during SARS-Cov-2^ORF3a^ expression. Cells were transfected with the GFP-SARS-Cov-2^ORF3a^ construct overnight. (E) Cell death analysis of supernatants of human peripheral blood monocyte-derived macrophages (hMDM) by a LDH release assay during hemozoin exposure. Cells were treated with 10µg/ml hemozoin for 4h in the presence or absence of 1 μg/ml cycloheximide (CHX). (F) Quantification using AMNIS of cell death by Live/Dead^TM^ stain kit in hMDM during silica treatment. Cells were treated with 200 µg/mL silica for 4h in the presence or absence of 1 μg/ml cycloheximide (CHX), and then stained using Live/Dead^TM^ stain kit (ThermoFisher). (G) Quantification using AMNIS of cell death by Live/Dead^TM^ stain kit in hMDM during the treatment of tau oligomer. Cells were treated with 10 µg/mL tau oligomer for 4h in the presence or absence of 1 μg/ml cycloheximide (CHX), and then stained using Live/Dead^TM^ stain kit (ThermoFisher). CTR, control. Data, means ± SEM (n = 3); HCM: n ≥ 3 (each experiment: 500 valid primary objects/cells per well, ≥5 wells/sample). *p < 0.05, **p < 0.01, ANOVA. See also Figure S6.

In addition, we examined other disease-associated agents that damage lysosomes. Hemozoin, a crystalline and insoluble material generated during malaria infection, circulates in phagocytes and peripheral tissues, contributing to immunopathological effects(Anyona et al., 2022; Coronado et al., 2014; Guerra et al., 2019; Moore et al., 2004; Schwarzer et al., 1992; Weissbuch et al., 2008). Treatment of human monocytic THP-1 with hemozoin induced lysosomal damage, which was monitored by ALIX puncta formation serving as a lysosomal repair marker(Jia et al., 2020c; Radulovic et al., 2018; Skowyra et al., 2018) (Fig. S6B). While a previous report showed that the acidification of lysosomes in hemozoin-fed monocytes was normal(Schwarzer et al., 2001), our finding suggest that hemozoin can perturb lysosomal membranes. This discrepancy in observations may depend on cell types, dosage and the duration of the treatment. In addition, the exposure to hemozoin resulted in both SG formation and eIF2α phosphorylation (Figs. S6C, D). Blocking SG formation with cycloheximide in hMDM cells, showed an increased cell death as measured by LDH release assay in response to hemozoin treatment (Fig. 7E). Moreover, we examined other lysosomal damaging agents, such as silica crystals associated with silicosis(Hornung et al., 2008; Mossman et al., 1998; J. Wang et al., 2017) and tau aggregates implicated in Alzheimer’s disease(Flavin et al., 2017; C. Papadopoulos et al., 2017). We have previously reported that both silica crystals and tau aggregates induce lysosomal damage, leading to SG formation(Jia et al., 2022). This effect was further confirmed by detecting eIF2α phosphorylation in hMDM cells in response to the treatment of silica crystals or tau aggregates (Fig. S6D). The prevention of SG formation with cycloheximide during the treatment of silica crystals or tau aggregates led to an augmented cell death as assessed using an AMNIS imaging flow cytometer in hMDM cells (Figs. 7F, G). Similarly, the application of another SG inhibitor, ISRIB to inhibit SG formation triggered by silica crystals or tau aggregate, produced a comparable effect on cell death of hMDM cells, measured by PI uptake assay (Figs. S6E, F). In summary, our finding suggest that SG formation induced by lysosomal damage is important for cell survival against diverse pathogenic challenges associated with major human diseases.

## DISCUSSION

In this study, we uncovered the regulation and significance of SG formation in response to lysosomal damage, providing insights into the interaction between membrane-bound organelles and membrane-less condensates. Through unbiased approaches including proteomic analysis and high content microscopy, we defined a novel signaling pathway that transmits calcium leakage from damaged lysosomes to induce eIF2α phosphorylation, ultimately leading to SG formation, thus promoting cell survival. This study aligns with recent research indicating the role of SGs in plugging damaged membrane and aiding in lysosomal repair(Bussi et al., 2023), underscoring SG formation as a vital cellular protective mechanism against lysosomal damage, essential for survival.

How does the cell detect lysosomal damage to initiate SGs? Our study revealed the significant involvement of a calcium signal in this process. Lysosomes function as key intracellular calcium reservoirs for various cellular activities(Lloyd-Evans et al., 2020; Xu et al., 2015). We found that ALIX and ALG2 sense calcium leakage from damaged lysosomes, leading to the activation of ALIX’s role in regulating PKR’s activity. This involves the control of the association between PKR and its activator PACT, resulting in the phosphorylation of eIF2α. Notably, we found that the role of ALIX and ALG2 in controlling eIF2α phosphorylation is distinct from their established function in ESCRT-mediated lysosomal repair. This suggests the multifaceted roles of ALIX and ALG2 as calcium sensors in coordinating cellular responses to lysosomal damage. Furthermore, our findings also indicate the intricate and adaptable nature of calcium signaling pathways in coordinating various cellular defense mechanisms against lysosomal damage. This extends beyond their involvement in TFEB nuclear translocation and phosphoinositide-mediates rapid lysosomal repair(Medina et al., 2015; Nakamura et al., 2020; Tan et al., 2022).

SGs consist of RNA-binding proteins and untranslated mRNA, both playing a crucial role in the process of phase separation(Millar et al., 2023). In addition to the calcium signal we reported here as a trigger for SG formation during lysosomal damage, a recent study suggests that a decrease in pH can also induce SG formation on damaged lysosomes(Bussi et al., 2023). This is in line with the reported role of pH in G3BP1-driven SG condensation(Guillén-Boixet et al., 2020). However, the latter report indicates that pH may not directly regulate the RNA-binding affinity of G3BP1 but instead influences protein-protein interactions. It is worth noting that these experiments were conducted in an in vitro system and in the presence of mRNA. Therefore, it raises the possibility that multiple mechanisms may collaborate to trigger SG formation by controlling protein-protein interaction or the accumulation of untranslated mRNA in response to lysosomal damage. To understand the signaling mechanism responsible for the accumulation of untranslated mRNA, our study suggests a calcium-dependent pathway that induces untranslated mRNA for SG formation by controlling eIF2α phosphorylation. Thus, both pH and calcium-dependent pathways can collaboratively contribute to SG formation during lysosomal damage. Moreover, considering the central role of lysosomes as the main degradation center for diverse cellular components, such as RNA(Lawrence et al., 2019), and the recognition of lysosomal damage can be sensed by various cellular machinery(Aits et al., 2015; Jia et al., 2022; Napolitano et al., 2016; Chrisovalantis Papadopoulos et al., 2017), the leakage of certain lysosomal contents or the activation of other lysosomal damage sensors may also contribute to the activation of PKR, eIF2α phosphorylation or the regulation of SG formation.

Phosphorylation of eIF2α is a key event in SG formation, as it causes the shutdown in global translation and the accumulation of untranslated mRNA, which triggers the phase separation, ultimately leading to SG formation (Ivanov et al., 2019; Riggs et al., 2020). However, there are instances of SG formation that occur independently of eIF2α phosphorylation, potentially regulated by the translation shutdown through the mTORC1 pathway(Emara et al., 2012; Fujimura et al., 2012). Nevertheless, this does not appear to be the case for SG formation in response to lysosomal damage. Our data indicate that upon lysosomal damage eIF2α phosphorylation is the primary driver for SG formation, though the impact of mTORC1 inactivation on translation shutdown and SG formation cannot be entirely ruled out. Importantly, the uncoupled relationship between mTORC1 inactivation and eIF2α phosphorylation in SG formation may be attributed to their differential impacts on protein translation events and mRNA entry into SGs. For example, mTORC1 inactivation primarily inhibits the translation pre-initiation, while eIF2α phosphorylation can impede the recruitment of the large ribosomal subunit to mRNA(Holz et al., 2005; Jackson et al., 2010). Recent research suggests that having just one large ribosomal subunit on mRNA is enough to prevent the recruitment of mRNA into SGs, while extended ribosome-free regions on mRNA are insufficient for SG formation(Fedorovskiy et al., 2023). Thus, mTORC1 inactivation may result in extended ribosome-free regions on mRNA, but alone, it is insufficient to prompt mRNA entry into SGs. The prevention of large ribosomal subunits on mRNA through eIF2α phosphorylation appears to be a crucial factor triggering this process and contributing to SG formation in the context of lysosomal damage. In addition, through the examination of SG formation in galectin knockout cells, previously we reported(Jia et al., 2022) that galectin-8 does not influence SG formation. This finding supports that eIF2α phosphorylation and mTORC1 inactivation are dissociated events during lysosomal damage, as we have previously reported that galectin-8 can modulate mTORC1 activity under similar conditions(Jia et al., 2018). Furthermore, recent research has highlighted lysosomes as pivotal hubs in metabolic signaling, involving mTORC1 and AMPK pathways(Carroll et al., 2017; Jia et al., 2018; Jia et al., 2020b; Zoncu et al., 2011). Our findings regarding the regulation of eIF2α phosphorylation on damaged lysosomes, along with our earlier findings showing mTORC1 inactivation on damaged lysosomes(Jia et al., 2018), propose an innovative role for lysosomes as central command centers in orchestrating protein translation signaling during stress conditions.

Understanding the function of SGs during stress, especially in the context of lysosomal damage, remains limited. A recent report highlights the reparative role of SGs through their association with damaged lysosomes(Bussi et al., 2023). This finding aligns with our prior research; however in our study, we observed SGs at a distance from damaged lysosomes(Jia et al., 2022). This observation challenges the notion of SGs primarily serving as plugs and suggests a broader spectrum roles of SGs in response to lysosomal damage. Given the significance of SG formation in supporting cell survival during lysosomal damage, as reported here, it is highly unlikely that SGs can undertake multiple tasks in restoring cellular homeostasis for survival. For example, considering SGs sequester non-translating mRNA(Khong et al., 2017), they may play roles in protecting mRNA and controlling mRNA fate and the transcriptome during lysosomal damage. Moreover, SG formation intersects with the integrated stress response (ISR), which can optimize cell response by reprogramming gene expressions to promote cellular recovery(Pakos-Zebrucka et al., 2016). The impact of SG formation on ISR may also enhance cellular fitness. Additionally, the involvement of SGs in various cellular processes, e.g., intracellular transport dynamics, ribosome biogenesis and cell signaling (Gorsheneva et al., 2024; Ripin et al., 2023; M.L. Zhang et al., 2024), may further contribute to cellular survival upon lysosomal damage.

Recognizing lysosomal damage as a critical internal physiological trigger for SGs highlights the importance of enhancing our understanding of SG formation in disease contexts. We detected the role of SG formation in cell survival within disease-specific contexts using a series of pathological reagents to induce lysosomal damage. Given the strong association of these reagents with both lysosomal damage and SG formation, delving into the molecular mechanisms governing the interaction between lysosomal damage and SGs may provide valuable insights for future therapeutic efforts.

## Acknowledgments

We thank Dr. P. Ivanov (Brigham and Women’s Hospital and Harvard Medical School, Boston, MA) for U2OS wildtype and G3BP1&2 double knockout cells. We thank Drs. J. Rajaiya, K. Bhaskar and D. Perkins (University of New Mexico Health Sciences Center, Albuquerque, NM) for human adenovirus, tau aggregates and hemozoin and hMDM respectively.

This work was supported by a NIH AIM COBRE grant (P20GM121176) to J. Jia. Mass spectrometry analysis was supported by a NIH S10 grant (S10OD026918-01A1) to B. Phinney.

## Authors contributions

J. Duran, S. Poolsup and J. Jia conceptualized the study, designed the experiments and analyzed the data; J. Duran, S. Poolsup and J. Jia performed majority of the experiments; M. Salemi, B. Phinney, J. Pu and L. Allers performed and interpreted mass spectrometry data; M. Lemus performed HDOCK analyses; Q. Cheng performed hemozoin and hMDM. J. Jia supervised the project. J. Jia wrote the manuscript with input from J. Duran and S. Poolsup.

## Declaration of Interests

The authors declare no competing interests.

## MATERIALS AND METHODS

### Antibodies and reagents

Antibodies from Cell Signaling Technology were Phospho-eIF2α (Ser51)(1:1000 for WB), eIF2α (1:1000 for WB), Phospho-p70 S6 Kinase (Thr389)(108D2)(1:1000 for WB), 4EBP1(1:1000 for WB), Phospho-4EBP1(Ser65)(1:1000 for WB), PKR (1:1000 for WB), PACT (D9N6J)(1:1000 for WB), Myc (9B11)(1:1000 for WB), mTOR (7C10)(1:1000 for WB; 1:400 for IF), ATF4(D4B8)(1:1000 for WB) and G3BP2 (1:1000 for WB). Antibodies from Abcam were GFP (ab290)(for immunoprecipitation (IP) or 1:1000 for WB), CHMP4B(ab105767)(1:1000 for WB), TSG101(4A10)(ab83)(1:1000 for WB), PKR (phospho T446) (E120)(ab32036) (1:1000 for WB). Antibodies from Sigma Aldrich: FLAG M2 (F1804)(for IP and 1:1000 for WB). Antibodies from Proteintech: EIF4G1 (15704-1-AP) (1:200 for IF), CHMP2A (10477-1-AP) (1:500 for WB), MARK2 (15492-1-AP) (1:500 for WB) and ALG2 (15092-1-AP) (1:500 for WB). Antibodies from BioLegend: Galectin-3 (1:1000 for WB; 1:500 for IF) and ALIX (1:200 for IF). G3BP1(PA5-29455, 1:1000 for WB, 1:200 for IF), Alexa Fluor 488, 568, 647 (1:500 for IF) and secondary antibodies from ThermoFisher Scientific. Other antibodies used in this study were from the following sources: beta-Actin (C4)(1:1000 for WB) from Santa Cruz Biotechnology; LAMP2 (H4B4)(1:500 for IF) from DSHB of University of Iowa.

Reagents from Sigma Aldrich were Leu-Leu-methyl ester hydrobromide (LLOMe), Sodium(meta)arsenite, Puromycin dihydrochloride, Imidazolo-oxindole PKR inhibitor C16, ISRIB and cycloheximide. Reagents from ThermoFisher were Hoechst 33342, BAPTA-AM, LIVE/DEAD™ Fixable Green Dead Cell Stain Kit, Lipofectamine RNAiMAX Transfection Reagent, BP/LR Clonase Plus Enzyme Mix, Prolong Gold Antifade Mountant with DAPI, Human M-CSF Recombinant Protein, Propidium Iodide (PI) solution, DMEM, Opti-MEM Reduced Serum Media, EBSS, PBS, Penicillin-Streptomycin, Fetal Bovine Serum, NP40 Cell Lysis Buffer, Anti-HA Magnetic Beads, Dynabeads Protein G, Streptavidin Magnetic Beads and LysoTracker Red DND-99. Reagents from Promega were CytoTox 96® Non-Radioactive Cytotoxicity Assay, FuGENE HD Transfection Reagent and ProFection Mammalian Transfection System. Other reagents used in this study were from the following sources: 5’-Cy3-Oligo d(T)30 from GeneLink (26-4330-02); Silica crystal from US Silica (MIN-U-SIL-15); Protease Inhibitor from Roche (11697498001). Wildtype human adenovirus species C2 (HAdV-C2^WT^) and its protease-deficient mutant TS1 (HAdV-C2^TS1^) were supported by Dr. Jaya Rajaiya (University of New Mexico Health Sciences Center, Albuquerque, NM). Hemozoin was supported by Dr. Douglas Perkins (University of New Mexico Health Sciences Center, Albuquerque, NM). Tau aggregates were supported by Dr. Kiran Bhaskar (University of New Mexico Health Sciences Center, Albuquerque, NM).

### Cells and cell lines

U2OS, HEK293T and THP-1 cells were from ATCC. Human peripheral blood monocyte-derived macrophages (hMDM) were derived from peripheral blood mononuclear cells (PBMCs) obtained from UNM Center for Global Health, details below. Cell lines for LysoIP were generated using constructs obtained from Addgene, details below. Knockout cell lines were generated by CRISPR/Cas9-mediated knockout system, and knockdown cell lines were generated by small interfering RNAs (siRNAs) from GE Dharmacon (siGENOME SMART pool), details below. U2OS G3BP1-GFP cell line was generated using Flp-In system (ThermoFisher), details below. U2OS wildtype (WT) and G3BP1&2 double knockout (ΔΔG3BP1/2) cells were from Dr. Pavel Ivanov (Brigham and Women’s Hospital and Harvard Medical School, Boston, MA).

### Cultured human peripheral blood monocyte-derived macrophages

A 40-50 mL blood draw was collected from a healthy, consenting adult volunteer. Keeping different donors separated, blood in 10 mL vacutainers was pooled into two 50 mL conical tubes and the volume was brought to 50 mL with sterile 1 X PBS followed by mixing inversely. 25 mL of the blood mix were carefully layered onto 20 mL of Ficoll (Sigma, #1077) in separate conical tubes and centrifuged at 2000 rpm for 30 min at 22 °C. The buffer layer containing human peripheral blood monocytes (PBMCs) was removed, pooled, washed with 1X PBS twice and resuspended in 20 mL RPMI media with 10 % human AB serum and Primocin. PBMCs were cultured in RPMI 1640 with GlutaMAX and HEPES (Gibco), 20 % FBS and 200 ng/mL Human M-CSF Recombinant Protein (ThermoFisher). Six days after the initial isolation, differentiated macrophages were detached in 0.25 % Trypsin-EDTA (Gibco) and seeded for experiments.

### Plasmids, siRNAs, and transfection

Plasmids used in this study, e.g., eIF2α, ALIX, PKR, and PACT cloned into pDONR221 using BP cloning, and expression vectors were made utilizing LR cloning (Gateway, ThermoFisher) in appropriate pDEST vectors for immunoprecipitation assay. Small interfering RNAs (siRNAs) were from Horizon Discovery (siGENOME SMART pool). Plasmid transfections were performed using the ProFection Mammalian Transfection System, FuGENE® HD Transfection Reagent (Promega), or Lipofectamine 2000 Transfection Reagent (ThermoFisher). siRNAs were delivered into cells using Lipofectamine RNAiMAX (ThermoFisher).

### Generation of CRISPR mutant cells

PKR knockout cells were generated by CRISPR/Cas9-mediated knockout system. The lentiviral vector lentiCRISPRv2 carrying both Cas9 enzyme and a gRNA transfected into HEK293T cells together with the packaging plasmids psPAX2 and pCMV-VSV-G (Addgene) at the ratio of 5:3:2. PKR: gRNA1: GATGGAAGAGAATTTCCAGA; gRNA2: AGTGTGCATCGGGGGTGCAT; gRNA3: TGGTACAGGTTCTACTAAAC (ABM, 19075111). Two days after transfection, the supernatant containing lentiviruses was collected. Cells were infected by the mixed lentiviruses containing gRNA1-3. 36 h after infection, the cells were selected with puromycin (2 µg/mL) for one week in order to select knockout cells. Knockout cells were confirmed by western blot. Selection of single clones was performed by dilution in 96-well, which were confirmed by western blots.

### Generating G3BP1-GFP cell line

Transfected U2OS Flp-In cells (generated by Flp-In system, ThermoFisher) with G3BP1-GFP reconstructed plasmid and the pOG44 expression plasmid at ration of 9:1. 24 h after transfection, washed the cells and added fresh medium to the cells. 48 h after transfection, split the cells into fresh medium around 25 % confluent. Incubate the cells at 37 °C for 2-3 h until they have attached to the culture dish. Then the medium was removed and added with fresh medium containing 100 µg/mL hygromycin. Cells were further fed with selective medium every 3-4 days until single cell clone can be identified. Picked hygromycin-resistant clones and expanded each clone to test.

### LysoIP assay

Lentiviruses constructs for generating stable LysoIP cells were purchased from Addgene. HEK293T cells were transfected with pLJC5-TMEM192-3xHA or pLJC5-TMEM192-2xFLAG constructs in combination with psPAX2 and pCMV-VSV-G packaging plasmids, at the ratio of 5:3:2, 60 h after transfection, the supernatant containing lentiviruses was collected and centrifuged to remove cells and then frozen at −80 °C. To establish LysoIP stably expressing cell lines, cells were plated in 10cm dish in DMEM with 10 % FBS and infected with 500 μL of virus-containing media overnight, then add puromycin for selection.

Selected cells in 15 cm plates with 90 % confluency were used for each LysoIP. Cells with or without treatment were quickly rinsed twice with PBS and then scraped in 1mL of KPBS (136 mM KCl, 10 mM KH_2_PO_4_, pH7.25 was adjusted with KOH) and centrifuged at 3000 rpm for 2 min at 4 °C. Pelleted cells were resuspended in 950 μL KPBS and reserved 25 μL for further processing of the whole-cell lysate. The remaining cells were gently homogenized with 20 strokes of a 2 mL homogenizer. The homogenate was then centrifuged at 3000 rpm for 2 min at 4 °C and the supernatant was incubated with 100 μL of KPBS prewashed anti-HA magnetic beads (ThermoFisher) on a gentle rotator shaker for 15 min. Immunoprecipitants were then gently washed three times with KPBS and eluted with 2 x Laemmli sample buffer (Bio-Rad) and subjected to immunoblot analysis.

### High content microscopy (HCM) analysis

Cells in 96 well plates were fixed in 4 % paraformaldehyde for 5 min. Cells were then permeabilized with 0.1 % saponin in 3 % Bovine serum albumin (BSA) for 30 min followed by incubation with primary antibodies for 2 h and secondary antibodies for 1 h. The analysis of Poly(A) RNA involved diluting a stock of 5’-labeled Cy3-Oligo-dT(30) stock (GeneLink) to a final concentration of 1 ng/μL, and incubation at 37 °C for at least one hour. Hoechst 33342 staining was performed for 3 min. HCM with automated image acquisition and quantification was carried out using a Cellomics HCS scanner and iDEV software (ThermoFisher). Automated epifluorescence image collection was performed for a minimum of 500 cells per well.

Epifluorescence images were machine analyzed using preset scanning parameters and object mask definitions. Hoechst 33342 staining was used for autofocus and to automatically define cellular outlines based on background staining of the cytoplasm. Primary objects were cells, and regions of interest (ROI) or targets were algorithm-defined by shape/segmentation, maximum/minimum average intensity, total area and total intensity, to automatically identify puncta or other profiles within valid primary objects. All data collection, processing (object, ROI, and target mask assignments) and analyses were computer driven independently of human operators. HCM provides a variable statistic since it does not rely on parametric reporting cells as positive or negative for a certain marker above or below a puncta number threshold.

### PI uptake assay

20,000 cells were plated in each well of a 96-well plate. Subsequently, cells were treated with lysosomal damaging agents, such as LLOMe. PI (propidium iodide) uptake was measured after 5 min incubation with 100 μg/mL diluted PI solution (ThermoFisher) in complete medium at 37 °C. After PI incubation, cells were fixed with 4% paraformaldehyde and stained with Hoechst 33342 for HCM analysis.

### LDH release assay

Each well of a 96-well plate was initially plated with 20,000 cells. Cells were treated with lysosomal damaging agents as indicated. Following this, the supernatant was measured for LDH (Lactate dehydrogenase) release using the kit of CytoTox 96® Non-Radioactive Cytotoxicity Assay (Promega, G1780), according to the manufacturer’s instructions.

### Amnis flow cytometry analysis

Cells after treatment were washed with 3% BSA in PBS supplemented with 0.1% of NaN3 before staining. Cells were stained using LIVE/DEAD™ Fixable Green Dead Cell Stain Kit (ThermoFisher) following the manufacturer’s instructions. After staining, cells were then resuspended with 3% BSA in PBS supplemented with 0.1% of NaN3 until acquisition on Amins ImageStreamx MKII (ISx, EMD Millipore, Seattle, WA, USA).

### LysoTracker assay

LysoTracker (LTR) Staining Solution was prepared by freshly diluting 2 μL of LTR stock solution (1 mM LysoTracker Red DND-99; Sigma Aldrich, L7528) in 1 mL of medium. 10 μL of Lyso-Tracker Staining Solution was added to 90 μL of medium each well in 96 well plates (final volume 100 μL per well, final concentration 0.2 μM LTR) and adherent cells incubated at 37°C for 30 min protected from light. Wells were rinsed gently by 1 × PBS and fixed in 4% paraformaldehyde for 2 min. Wells were washed once in 1 × PBS and nuclei stained with Hoechst 33342 for 2 min before analyzing the plates by HCM.

### Co-immunoprecipitation assay

Cells transfected with 8-10 μg of plasmids were lysed in NP-40 buffer (ThermoFisher) supplemented with protease inhibitor cocktail (Roche, 11697498001) and 1 mM PMSF (Sigma, 93482) for 30 min on ice. Supernatants were incubated with (2-3 μg) antibodies overnight at 4 °C. The immune complexes were captured with Dynabeads (ThermoFisher), followed by three times washing with 1 x PBS. Proteins bound to Dynabeads were eluted with 2 x Laemmli sample buffer (Bio-Rad) and subjected to immunoblot analysis. Immunoblotting images were visualized and analyzed using ImageLab v.6.0.0.

### Immunofluorescence confocal microscopy analysis

Cells were plated onto coverslips in 6-well plates. After treatment, cells were fixed in 4 % paraformaldehyde for 5 min followed by permeabilization with 0.1 % saponin in 3 % BSA for 30 min. Cells were then incubated with primary antibodies for 2 h and appropriate secondary antibodies Alexa Fluor 488 or 568 (ThermoFisher) for 1 h at room temperature. Coverslips were mounted using Prolong Gold Antifade Mountant (ThermoFisher). Images were acquired using a confocal microscope (META; Carl Zeiss) equipped with a 63 3/1.4 NA oil objective, camera (LSM META; Carl Zeiss), and AIM software (Carl Zeiss).

### APEX2-Labeling and streptavidin enrichment for LC/MS/MS DIA analysis

HEK293T cells transfected APEX2 - eIF2α were incubated with 1 mM LLOMe for 1 h (confluence of cells remained at 70-80 %). Cells were next incubated in 500 mM biotin-phenol (AdipoGen) for the last 45 min of LLOMe incubation. A 1 min pulse with 1mM H_2_O_2_ at room temperature was stopped with quenching buffer (10 mM sodium ascorbate, 10 mM sodium azide and 5 mM Trolox in Dulbecco’s Phosphate Buffered Saline (DPBS)). All samples were washed twice with quenching buffer, and twice with DPBS.

For mass spectrometry analysis, cell pellets were lysed in 500 mL ice-cold lysis buffer (6 M urea, 0.3 M Nacl, 1 mM EDTA, 1 mM EGTA, 10 mM sodium ascorbate, 10 mM sodium azide, 5 mM Trolox, 1%glycerol and 25 mm Tris/HCl, PH 7.5) for 30 min by gentle pipetting. Lysates were clarified by centrifugation and protein concentrations determined as above. Streptavidin– coated magnetic beads (Pierce) were washed with lysis buffer. 3 mg of each sample was mixed with 100 mL of streptavidin bead. The suspensions were gently rotated at 4 °C for overnight to bind biotinylated proteins. The flowthrough after enrichment was removed and the beads were washed in sequence with 1 mL IP buffer (150 mM NaCl, 10 mM Tris-HCl pH8.0, 1 mM EDTA, 1 mM EGTA, 1 % Triton X-100) twice; 1 mL 1M KCl; 1mL of 50 mM Na_2_CO_3_; 1 mL 2M Urea in 20 mM Tris HCl pH8; 1 mL IP buffer. Biotinylated proteins were eluted, 10% of the sample processed for Western Blot and 90% of the sample processed for LC/MS/MS DIA (data-independent acquisition mass spectrometry) analysis.

LC/MS/MS DIA were performed at UC Davis Proteomics Core Facility (Davis, CA). Protein samples on magnetic beads were washed four times with 200 µL of 50 mM triethyl ammonium bicarbonate (TEAB) with a 20 min shake time at 4°C in between each wash. Roughly 2.5 mg of trypsin was added to the bead and TEAB mixture and the samples were digested over night at 800 rpm shake speed. After overnight digestion the supernatant was removed, and the beads were washed once with enough 50 mM ammonium bicarbonate to cover. After 20 min at a gentle shake the wash is removed and combined with the initial supernatant. The peptide extracts are reduced in volume by vacuum centrifugation and a small portion of the extract is used for fluorometric peptide quantification (ThermoFisher). One microgram of sample based on the fluorometric peptide assay was loaded for each LC/MS/MS analysis.

Peptides were separated on an Easy-spray 100 mm x 25 cm C18 column using a Dionex Ultimate 3000 nUPLC. Solvent A=0.1 % formic acid, Solvent B=100 % Acetonitrile 0.1 % formic acid. Gradient conditions = 2 % B to 50 % B over 60 minutes, followed by a 50 %-99 % B in 6 min and then held for 3 min than 99 % B to 2 % B in 2 min. Total Run time = 90 min. Thermo Scientific Fusion Lumos mass spectrometer running in data independent analysis (DIA) mode. Two gas phases fractionated (GFP) injections were made per sample using sequential 4 Da isolation widows. GFP1 = m/z 362-758, GFP 2 = m/z 758-1158. Tandem mass spectra were acquired using a collision energy of 30, resolution of 30K, maximum inject time of 54 ms and a AGC target of 50K.

### DIA Quantification and Statistical Analysis

DIA data was analyzed using Spectronaut. Raw data files were converted to mzML format using ProteoWizard (3.0.11748). Analytic samples were aligned based on retention times and individually searched against Pan human library http://www.swathatlas.org/ with a peptide mass tolerance of 10.0 ppm and a fragment mass tolerance of 10.0 ppm. Variable modifications considered were: Modification on M M and Modification on C C. The digestion enzyme was assumed to be Trypsin with a maximum of 1 missed cleavage site(s) allowed. Only peptides with charges in the range <2..3> and length in the range <6..30> were considered. Peptides identified in each sample were filtered by Percolator (3.01.nightly-13-655e4c7-dirty) to achieve a maximum FDR of 0.01. Individual search results were combined and peptide identifications were assigned posterior error probabilities and filtered to an FDR threshold of 0.01 by Percolator (3.01.nightly-13-655e4c7-dirty). Peptide quantification was performed by Encyclopedia (0.8.1). For each peptide, the 5 highest quality fragment ions were selected for quantitation. Proteins that contained similar peptides and could not be differentiated based on MS/MS analysis were grouped to satisfy the principles of parsimony. Proteins with a minimum of 2 identified peptides were thresholded to achieve a protein FDR threshold of 1.0%. Raw data and Spectronaut results are in Table S1.

### Quantification and statistical analysis

Data in this study are presented as means ± SEM (n ≥ 3). Data were analyzed with either analysis of variance (ANOVA) with Tukey’s HSD post-hoc test, or a two-tailed Student’s t test. For HCM, n ≥ 3 includes in each independent experiment: 500 valid primary objects/cells per well, from ≥ 5 wells per plate per sample. Statistical significance was defined as: † (not significant) p ≥ 0.05 and *p < 0.05, **p < 0.01.

### Data availability

Raw MS DIA data of APEX2 - eIF2α in HEK293T cells have been deposited at the MassIVE proteomics repository (MSV000093768) and Proteome Exchange (PXD048258).

---

**Figure S1.**
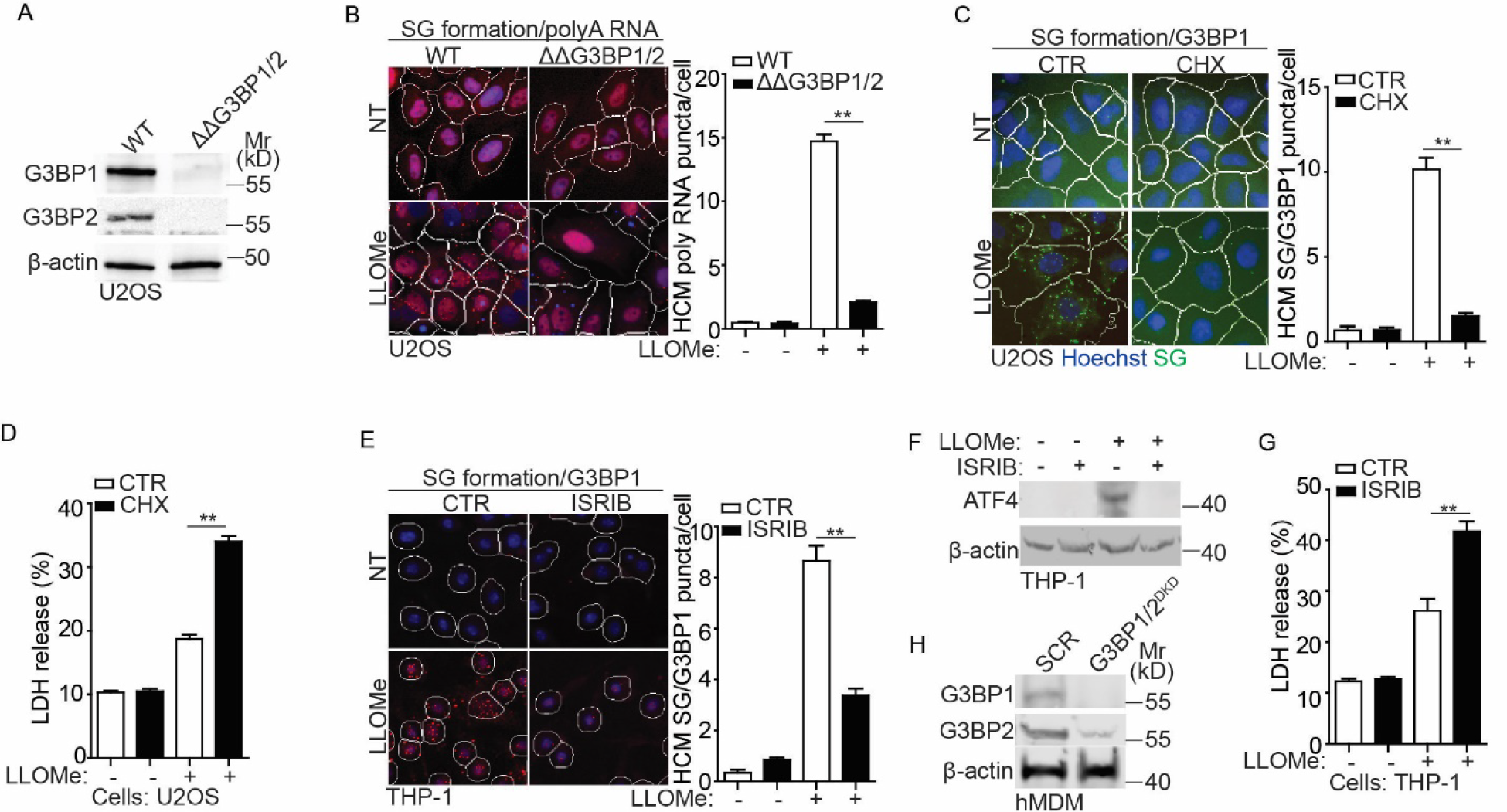
Stress granule formation is important for cell survival during lysosomal damage. (A) Immunoblot analysis of G3BP1 and G3BP2 in U2OS WT and ΔΔG3BP1/2 cells. (B) Quantification by high-content microscopy (HCM) of polyA RNA (Cy3-oligo[dT]) by FISH in U2OS WT and ΔΔG3BP1/2 cells. Cells were treated with 2 mM LLOMe for 30 min. White masks, algorithm-defined cell boundaries (primary objects); red masks, computer-identified polyA RNA puncta (target objects). (C) Quantification by HCM of G3BP1 puncta in U2OS cells. Cells were treated with 2 mM LLOMe in the presence or absence of 10 μg/ml cycloheximide (CHX) for 30 min. White masks, algorithm-defined cell boundaries; green masks, computer-identified G3BP1 puncta. (D) Cell death analysis of supernatants of U2OS cells by a LDH release assay. Cells were treated with 2 mM LLOMe in the presence or absence of 10 μg/ml CHX for 30 min. (E) Quantification by HCM of G3BP1 puncta in human monocytic THP-1 cells. Cells were treated with 1 mM LLOMe in the presence or absence of 200 nM ISRIB for 30 min. White masks, algorithm-defined cell boundaries; red masks, computer-identified G3BP1 puncta. (F) Immunoblot analysis of ATF4 in THP-1 cells treated with 1 mM LLOMe in the presence or absence of 200 nM ISRIB for 30 min. (G) Cell death analysis of supernatants of THP-1 cells by a LDH release assay. Cells were treated with 1 mM LLOMe in the presence or absence of 200 nM ISRIB for 30 min. (H) Immunoblot analysis of the protein level of G3BP1 and G3BP2 in hMDM transfected with scrambled siRNA as control (SCR) or G3BP1 and G3BP2 siRNA for double knockdown (DKD). CTR, control; NT, untreated cells. Data, means ± SEM (n = 3); HCM: n ≥ 3 (each experiment: 500 valid primary objects/cells per well, ≥5 wells/sample). **p < 0.01, ANOVA. See also Figure 1.

**Figure S2.**
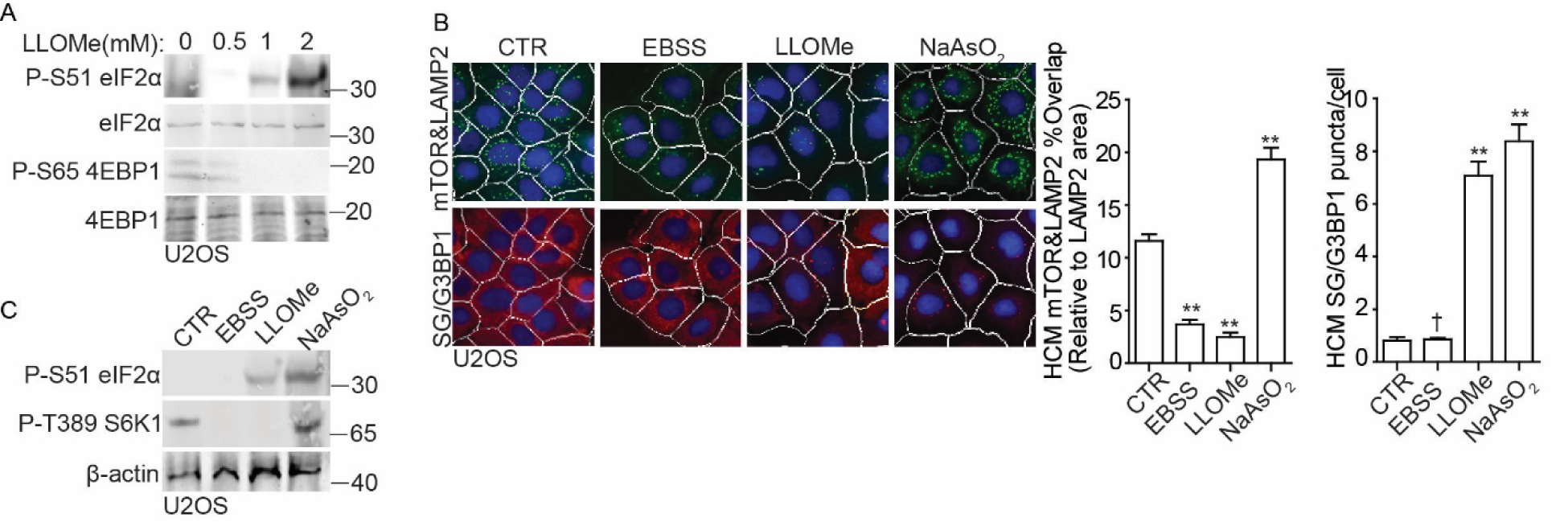
The eIF2α and mTORC1 signaling pathways are uncoupled in response to lysosomal damage. (A) Immunoblot analysis of phosphorylation of eIF2α (S51) and 4EBP1 (S65) in U2OS cells treated with the indicated dose of LLOMe for 30 min. (B) Quantification by HCM of overlaps between mTOR and LAMP2 or G3BP1 puncta in U2OS cells. Cells were treated with EBSS, 2 mM LLOMe or 100 µM NaAsO2 for 30 min. White masks, algorithm-defined cell boundaries; green masks, computer-identified overlap between mTOR and LAMP2; red masks, computer-identified G3BP1 puncta. (C) Immunoblot analysis of phosphorylation of eIF2α (S51) and S6K1 (T389) in U2OS cells treated as in (B). CTR, control. Data, means ± SEM (n = 3); HCM: n ≥ 3 (each experiment: 500 valid primary objects/cells per well, ≥5 wells/sample). † p ≥ 0.05 (not significant), **p < 0.01, ANOVA. See also Figure 2.

**Figure S3.**
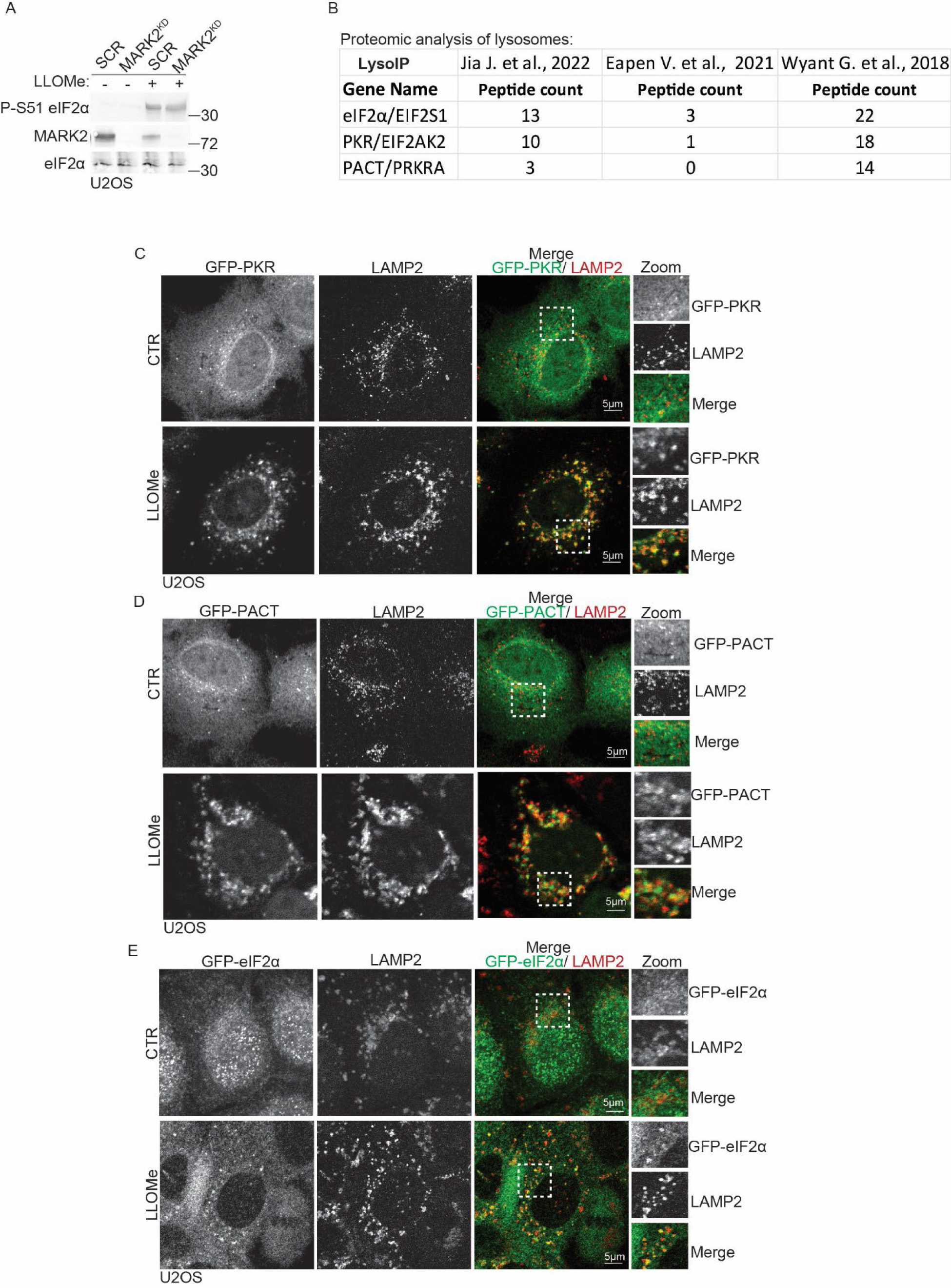
PKR, PACT and eIF2α are associated with damaged lysosomes. (A) Immunoblot analysis of phosphorylation of eIF2α (S51) in U2OS cells transfected with either scrambled siRNA as control (SCR) or MARK2 siRNA for knockdown (MARK2^KD^). Cells were treated with 2 mM LLOMe for 30 min. (B) Summary of the literature on the detected peptide count of PKR, PACT and eIF2α in the proteomic analysis of lysosomes based on LysoIP LC/MS/MS analysis. (C) Confocal microscopy imaging of GFP-PKR and LAMP2 in U2OS cells treated with 2 mM LLOMe for 30 min. Scale bar, 5 μm. (D) Confocal microscopy imaging of GFP-PACT and LAMP2 in U2OS cells treated with 2 mM LLOMe for 30 min. Scale bar, 5 μm. (E) Confocal microscopy imaging of GFP-eIF2α and LAMP2 in U2OS cells treated with 2 mM LLOMe for 30 min. Scale bar, 5 μm. See also Figure 3.

**Figure S4.**
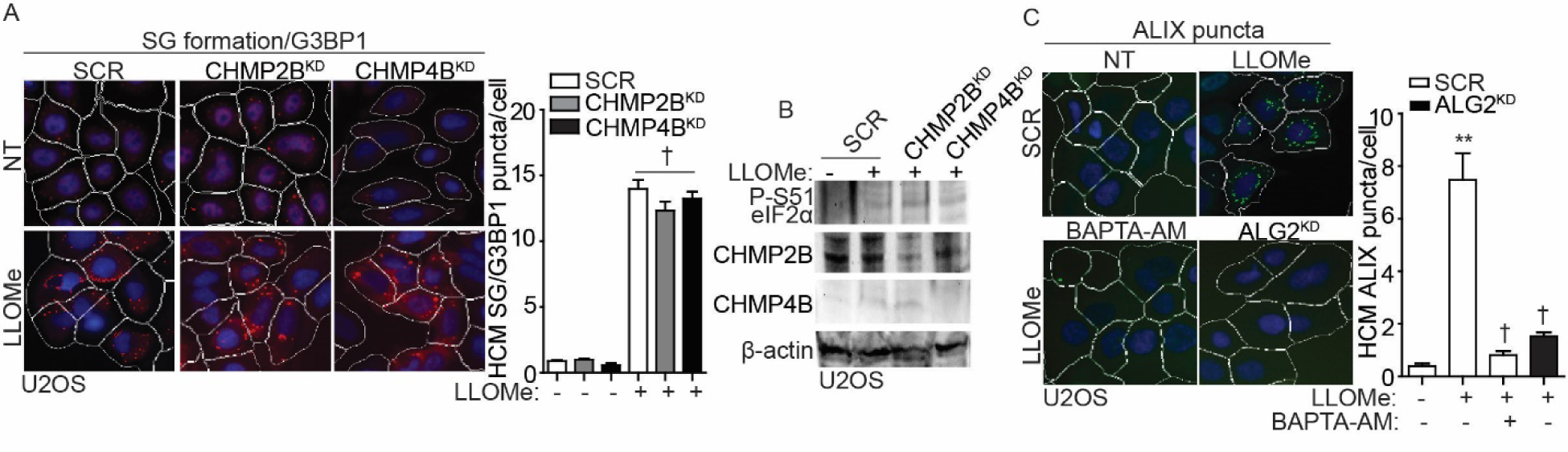
ALIX regulates stress granule formation during lysosomal damage. (A) Quantification by HCM of G3BP1 puncta in U2OS cells transfected with scrambled siRNA as control (SCR), CHMP2B siRNA for knockdown (CHMP2B^KD^) or CHMP4B siRNA for knockdown (CHMP4B^KD^). Cells were treated with 2 mM LLOMe for 30 min. White masks, algorithm-defined cell boundaries; red masks, computer-identified G3BP1 puncta. (B) Immunoblot analysis of phosphorylation of eIF2α (S51) in U2OS transfected with scrambled siRNA as control (SCR), CHMP2B siRNA for knockdown (CHMP2B^KD^) or CHMP4B siRNA for knockdown (CHMP4B^KD^), subjected to 2 mM LLOMe treatment for 30 min. (C) Quantification by HCM of ALIX puncta in U2OS cells transfected with scrambled siRNA as control (SCR), or ALG2 siRNA for knockdown (ALG2^KD^), or pre-treated with 15 µM BAPTA-AM for 1 h. Cells were treated with 2 mM LLOMe for 30 min. White masks, algorithm-defined cell boundaries; green masks, computer-identified ALIX puncta. NT, untreated cells. Data, means ± SEM (n = 3); HCM: n ≥ 3 (each experiment: 500 valid primary objects/cells per well, ≥5 wells/sample). † p ≥ 0.05 (not significant), **p < 0.01, ANOVA. See also Figure 4.

**Figure S5.**
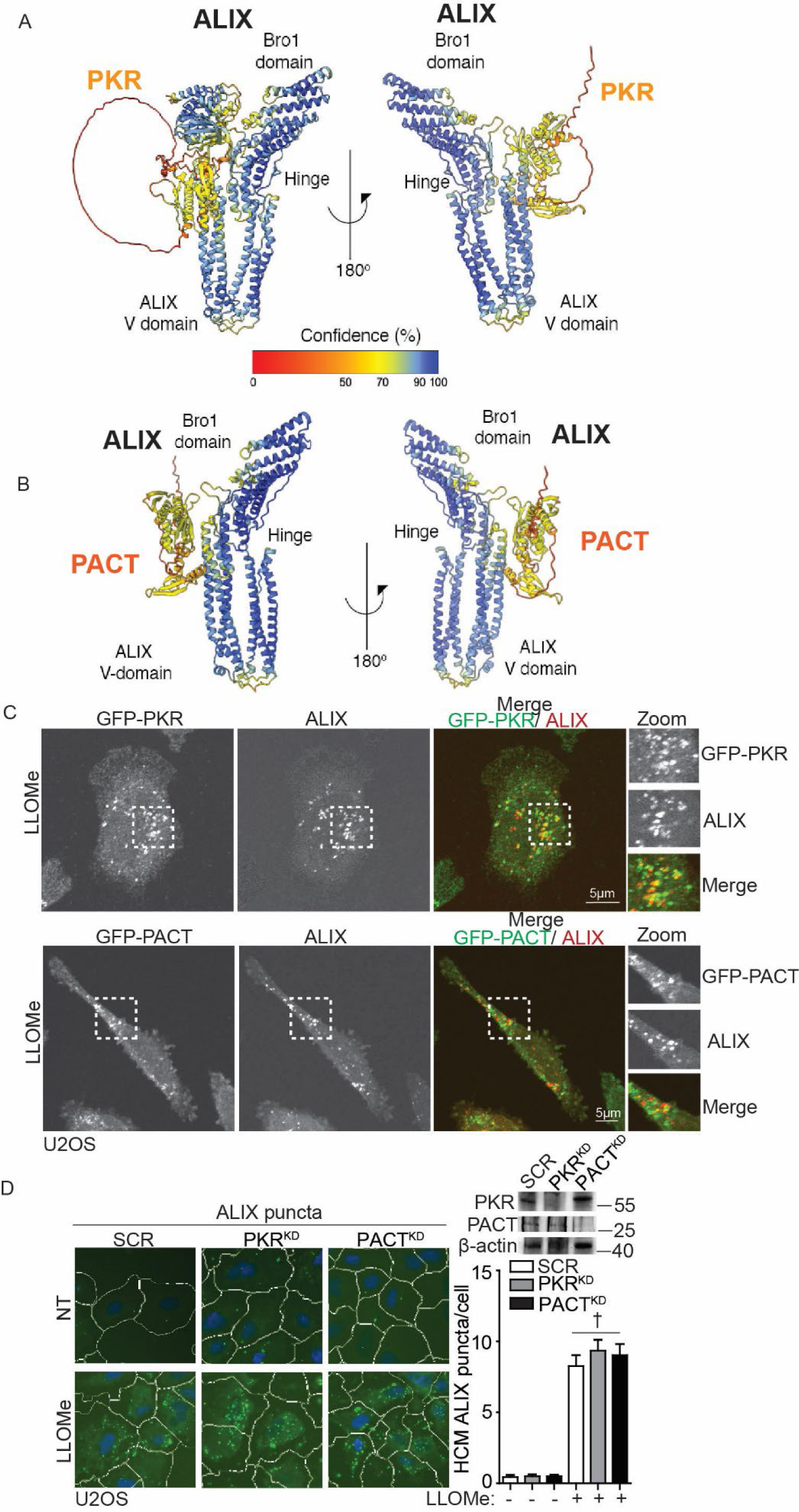
PKR and PACT associate with ALIX during lysosomal damage. (A) HDOCK-predicted interaction between ALIX and PKR. (B) HDOCK-predicted interaction between ALIX and PACT. (C) Confocal microscopy imaging of GFP-PKR/PACT and ALIX in U2OS cells treated with 2 mM LLOMe for 30 min. Scale bar, 5 μm. (D) Quantification by HCM of ALIX puncta in U2OS cells transfected with scrambled siRNA as control (SCR), PKR siRNA for knockdown (PKR^KD^), or PACT siRNA for knockdown (PACT^KD^). Cells were treated with 2 mM LLOMe for 30 min. White masks, algorithm-defined cell boundaries; green masks, computer-identified ALIX puncta. NT, untreated cells. Data, means ± SEM (n = 3); HCM: n ≥ 3 (each experiment: 500 valid primary objects/cells per well, ≥5 wells/sample). † p ≥ 0.05 (not significant), ANOVA. See also Figure 5.

**Figure S6.**
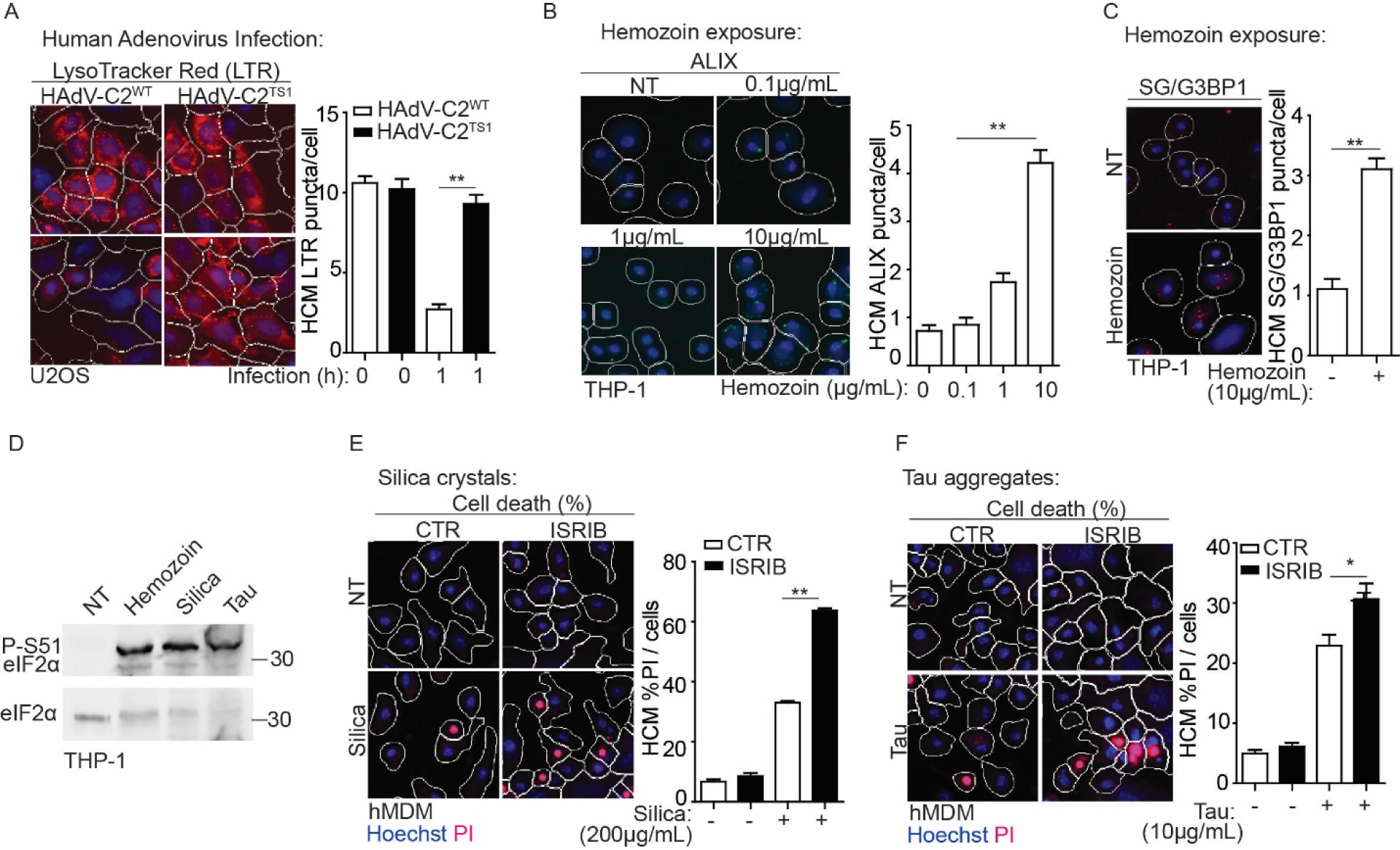
Stress granules are important for cell survival in response to lysosomal damage in disease states. (A) Quantification by HCM of status of acidified organelles assessed by LysoTracker Red (LTR) in U2OS cells infected with wildtype human adenovirus C2 (HAdV-C2^WT^) or C2 TS1 mutant (HAdV-C2^TS1^) at MOI=10 for 1h. White masks, algorithm-defined cell boundaries; red masks, computer-identified LTR puncta. (B) Quantification by HCM of ALIX puncta in THP-1 cells treated with hemozoin for 4h at the indicated dose. White masks, algorithm-defined cell boundaries; green masks, computer-identified ALIX puncta. (C) Quantification by HCM of G3BP1 puncta in THP-1 cells treated with 10 µg/ml hemozoin for 4h. White masks, algorithm-defined cell boundaries; red masks, computer-identified G3BP1 puncta. (D) Immunoblot analysis of phosphorylation of eIF2α (S51) in THP-1 cells treated with 10 µg/ml hemozoin, 200 µg/mL silica or 10 µg/mL tau oligomer for 4h. (E) Quantification by HCM of cell death by a propidium iodide (PI) uptake assay in human peripheral blood monocyte-derived macrophages (hMDM) during silica treatment. Cells were treated with 200 µg/mL silica for 4h in the presence or absence of 100 nM ISRIB, and then stained with propidium iodide PI (dead cells) and Hoechst-33342 (total cells). White masks, algorithm-defined cell boundaries; red masks, computer-identified PI+ nuclei. (F) Quantification by HCM of cell death by a propidium iodide (PI) uptake assay in human peripheral blood monocyte-derived macrophages (hMDM) during the treatment of tau oligomer. Cells were treated with 10 µg/mL tau oligomer for 4h in the presence or absence of 100 nM ISRIB, and then stained with propidium iodide PI (dead cells) and Hoechst-33342 (total cells). White masks, algorithm-defined cell boundaries; red masks, computer-identified PI+ nuclei. CTR, control; NT, untreated cells. Data, means ± SEM (n = 3); HCM: n ≥ 3 (each experiment: 500 valid primary objects/cells per well, ≥5 wells/sample). *p < 0.05, **p < 0.01, ANOVA. See also Figure 7.

## Notes

### Competing Interest Statement

The authors have declared no competing interest.

### Summary of Updates

This revised version of the manuscript has been updated to correct the author affiliation.

